# Absolute quantitative and base-resolution sequencing reveals comprehensive landscape of pseudouridine across the human transcriptome

**DOI:** 10.1101/2024.01.08.574649

**Authors:** Haiqi Xu, Linzhen Kong, Jingfei Cheng, Khatoun Al Moussawi, Xiufei Chen, Aleema Iqbal, Peter A. C. Wing, James M. Harris, Senko Tsukuda, Azman Embarc-Buh, Guifeng Wei, Alfredo Castello, Skirmantas Kriaucionis, Jane A. McKeating, Xin Lu, Chun-Xiao Song

**Affiliations:** Ludwig Institute for Cancer Research, Nuffield Department of Medicine, University of Oxford, Oxford OX3 7FZ, UK; Target Discovery Institute, Nuffield Department of Medicine, University of Oxford, Oxford OX3 7FZ, UK; Chinese Academy of Medical Sciences Oxford Institute, Nuffield Department of Medicine, University of Oxford, Oxford OX3 7BN, UK; Nuffield Department of Medicine, University of Oxford, Oxford OX3 7FZ, UK; MRC University of Glasgow Centre for Virus Research, Glasgow G61 1QH, UK; Department of Biochemistry, University of Oxford, Oxford OX1 3QU, UK

## Abstract

Pseudouridine (Ψ) is one of the most abundant modifications in cellular RNA. However, its function remains elusive, mainly due to the lack of highly sensitive and accurate detection methods. To address this challenge, we introduced 2-bromoacrylamide-assisted cyclization sequencing (BACS) for quantitative profiling of Ψ at single-base resolution. Based on novel bromoacrylamide cyclization chemistry, BACS enables a Ψ-to-C transition. Compared to previous methods, BACS allowed the precise identification of Ψ positions, especially in densely modified Ψ regions and consecutive uridine sequences. BACS successfully detected all known Ψ sites in human rRNA and spliceosomal snRNAs and generated the first quantitative Ψ map of human snoRNA and tRNA. Furthermore, BACS simultaneously detected adenosine-to-inosine (A-to-I) editing sites and *N*^1^-methyladenosine (m^1^A). Depletion of three key pseudouridine synthases (PUS) enabled us to elucidate the targets and sequence motifs of TRUB1, PUS7, and PUS1 in HeLa cells. We further applied BACS to Epstein-Barr virus (EBV)-encoded small RNAs (EBERs) and identified a highly abundant Ψ_114_ site in EBER2. Surprisingly, applying BACS to a panel of RNA viruses demonstrated the absence of Ψ in their viral transcripts or genomes, shedding light on differences in pseudouridylation between virus families. We anticipate BACS to serve as a powerful tool to uncover the biological importance of Ψ in future studies.

## Introduction

Pseudouridine (Ψ), as the C-C glycoside isomer of uridine, is the most abundant post-transcriptional modification in cellular RNA^1,2^. It is not only prevalent in nearly all kinds of non-coding RNAs (ncRNAs), including ribosomal RNA (rRNA), small nuclear RNA (snRNA), and transfer RNA (tRNA)^3^, but has also been known to be present in messenger RNA (mRNA)^4,5^. Ψ plays an important role in splicing, translation, RNA stability, and RNA-protein interactions^6^. In eukaryotes, Ψ is installed by various pseudouridine synthases (PUS)^7,8^, which have been shown to associate with many diseases including cancer^9–14^. Therefore, establishing an accurate and sensitive method to detect Ψ is highly desirable.

Traditionally, the detection of Ψ has been reliant on the *N*-cyclohexyl-*N*’-(2-morpholinoethyl)carbodiimide methyl-*p*-toluenesulfonate (CMC) chemistry^15^. CMC can react with amide or imide functional group in nucleobases (for example, amide for guanosine and imide for uridine)^16,17^, while it can form a more stable adduct with *N*^3^ of Ψ, therefore enabling discrimination of Ψ with uridine through subsequent alkaline treatment (pH 10.4)^18^. Since *N*^3^-CMC adduct of Ψ strongly interferes with base pairing and would lead to reverse transcription (RT) truncations, CMC chemistry has been widely applied to transcriptional-wide mapping of Ψ, as shown in Pseudo-seq^4^, Ψ-seq^5^, PSI-seq^19^, and CeU-seq^20^. However, CMC-based methods have low labeling efficiency and selectivity for Ψ, making it intrinsically difficult to distinguish between true Ψ signals and background noise emerging from other bases and RNA secondary structures^21,22^. Consequently, CMC-based methods lack stoichiometry information of Ψ.

Recently, RBS-seq reexamined the bisulfite (BS)-mediated conversion of Ψ^23,24^, and surprisingly found that the Ψ-BS adduct could lead to deletion signatures during RT, thus providing a novel way to detect Ψ (ref.^25,26^). Since unmodified cytosine would also be deaminated to uridine in conventional BS reaction^27,28^, BID-seq and similarly designed PRAISE further optimized BS treatment at near neutral pH to eliminate most of the side reaction on unmodified cytosine, enabling quantitative detection of Ψ (ref.^29,30^). Although BID-seq substantially improved this approach, it still suffered from low deletion rates in certain sequence contexts. Crucially, due to the deletion signature, BS-based methods cannot determine the exact position of Ψ in consecutive uridine sequences or consecutive Ψ sites and struggle to detect densely modified Ψ sites. Although local realignment can partially solve this issue, it can generate artefacts including overestimation of Ψ modification level and misidentification of sequence context^31^.

To overcome these limitations, we developed BACS for direct, quantitative, and base-resolution sequencing of Ψ based on new bromoacrylamide cyclization chemistry. BACS induces Ψ-to-C mutation rather than truncation or deletion signatures during RT, thereby providing higher resolution and more accurate quantification of Ψ stoichiometry compared with CMC- and BS-based methods. We applied BACS to various types of ncRNAs from HeLa, C666-1, Raji, and Elijah cells; and mRNA from HeLa cells to build a comprehensive map of Ψ across the human transcriptome. Besides Ψ mapping, BACS delivers simultaneous detection of adenosine-to-inosine (A-to-I) editing sites in mRNA and *N*^1^-methyladenosine (m^1^A) in tRNA. We further utilized BACS to elucidate genuine Ψ targets and sequence motifs of three key PUS enzymes (TRUB1, PUS7, and PUS1) in HeLa cells. Finally, we applied BACS to various RNA and DNA viruses to investigate the presence of Ψ in viral RNAs.

## Results

### Development of BACS

The most distinct difference between Ψ and uridine is the free *N*^1^ of Ψ, which is highly reactive towards Michael addition acceptors, such as acrylonitrile^32–36^, acrylamide^37^, and other acrylic compounds^38^. Selective labeling of Ψ by acrylonitrile has been widely used to distinguish Ψ from uridine in mass spectrometry^39^. However, a simple *N*^1^-adduct of Ψ with acrylic compounds would not induce mutation during RT. Further inspired by the formation and mutation profile of *O*^6^-methylguanine (inducing G-to-A mutation) and *O*^4^-methylthymine (inducing T-to-C mutation)^40,41^, we envisioned that installing an α-halogen group would induce tandem cyclization of the *N*^1^-acrylic adduct of Ψ through intramolecular *O*^2^-alkylation and finally lead to Ψ-to-C mutation^42^ (Fig. 1a). We initially tested this chemistry on a short Ψ-containing oligonucleotide with 2-bromoacrylamide and analyzed the reaction product by matrix-assisted laser desorption/ionization mass spectrometry (MALDI). We found a 69 Da increase of mass values, indicating the formation of a cyclized product (carbamido-1, *O*^2^-ethano Ψ, nce^1,2^Ψ) (Fig. 1b and Supplementary Fig. 1a). This reaction was further confirmed by ultra-high-performance liquid chromatography-tandem mass spectrometry (UHPLC-MS/MS) (Supplementary Fig. 1b). To validate the mutation signature of nce^1,2^Ψ, we applied BACS to a 72mer model RNA which contains two Ψ sites. Through RT and next-generation sequencing, we obtained 86.6% U-to-C mutation rates on these two sites, while U-to-R (R = A or G) mutation rates were lower than 1% (Supplementary Fig. 1c). Therefore, we confirm that the U-to-C mutation rate can serve as the conversion rate of BACS.

**Fig. 1.**
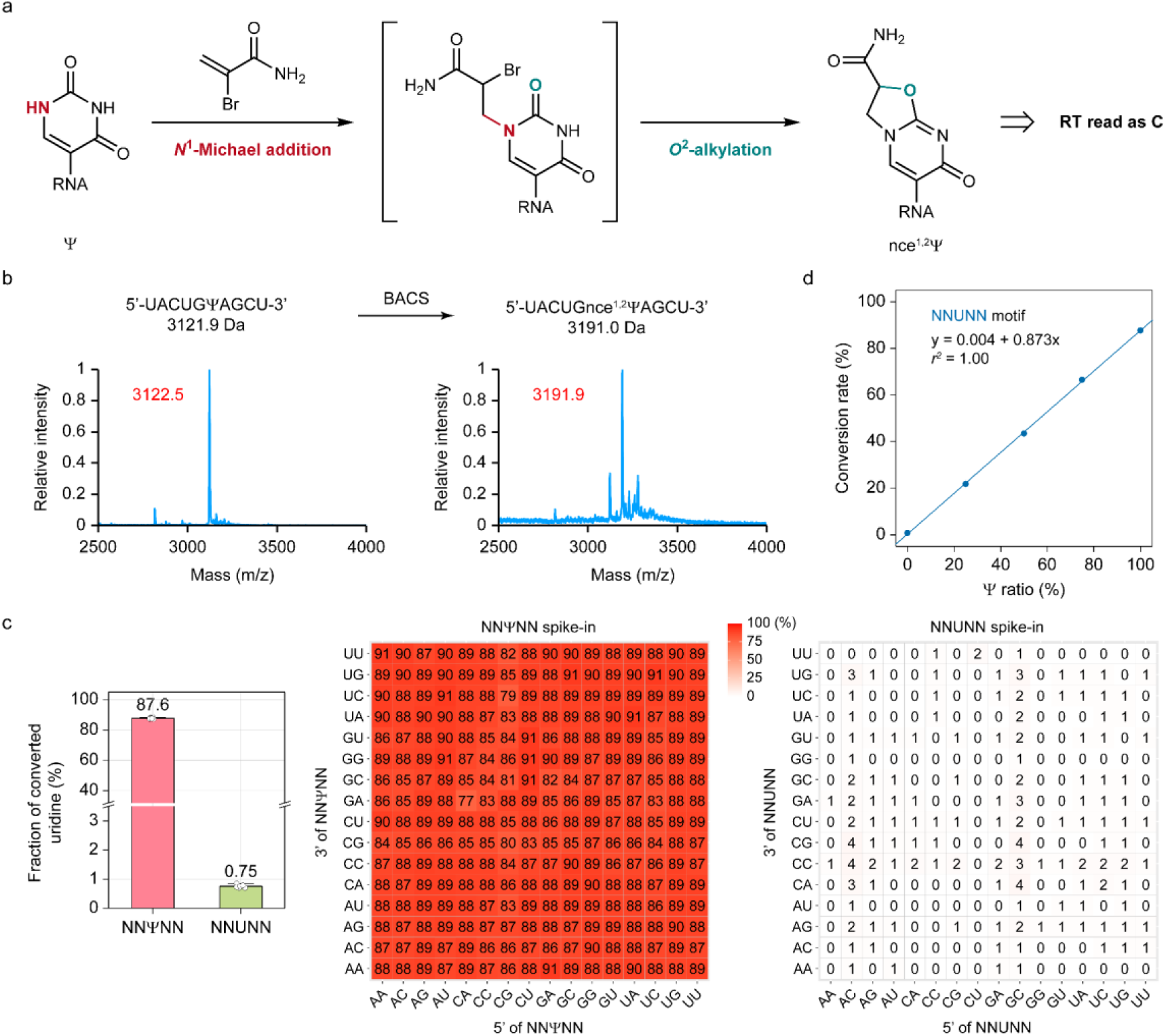
BACS achieved quantitative detection of Ψ through novel cyclization chemistry. **a.** Schematic overview of BACS reaction. **b.** MALDI characterization of BACS labeling of a 10mer Ψ-containing RNA oligonucleotide. Calculated mass is shown in black. Observed mass is shown in red. Experiment was performed once. **c.** Cumulative (left) and motif-dependent (middle and right) results of BACS conversion rates and false-positive rates on synthetic 30mer NNΨNN and NNUNN spike-in. Data are shown as means ± s.d. of six independent experiments (*n* = 6). **d.** BACS calibration curve for quantification of Ψ stoichiometry in NNUNN motif. Experiment was performed once.

To understand the sequence preference of BACS, we built libraries with synthetic 30mer RNA spike-in containing NNΨNN and NNUNN (N = A, C, G or U), respectively (Fig. 1c). After BACS, we observed an 87.6% conversion rate of Ψ and a 0.75% false-positive rate of uridine when accumulating all motifs. Among all the 256 motifs, 230 showed a conversion rate higher than 85% and 254 displayed a conversion rate higher than 80%, suggesting the high efficiency of BACS chemistry. We observed a low false-positive rate (<1%) in most motifs (213 out of 256 motifs). Certain motifs, especially those with one or more cytidines 5’- or 3’-flanking to the uridine site (for example, GCUCC and ACUCC), displayed slightly higher false-positive rates (3–4%), possibly due to the preferences of RT. Nevertheless, BACS clearly showed higher conversion rates than BID-seq both in general and in specific motifs^29^. By mixing NNΨNN and NNUNN spike-in in different ratios, we generated excellent linear calibration curves for accurate quantification of Ψ modification levels (*r*^2^ = 1.00, Fig. 1d and Supplementary Fig. 1d).

### Validation of BACS on human rRNA

To evaluate the performance of BACS, we applied it to cytosolic rRNA (cy-rRNA) from cervical and nasopharyngeal cancer cell lines HeLa and C666-1, respectively (Fig. 2a). Cy-rRNA is known to possess a series of highly conserved Ψ sites^43^. Using a 5% modification level cutoff, we detected 2, 40, and 62 Ψ sites in HeLa 5.8S, 18S, and 28S rRNAs, respectively (Fig. 2b). Most of the sites displayed a high level of Ψ modification (>80%), consistent with reports that Ψ sites are highly modified in human cy-rRNA^44^ (Fig. 2c–e and Supplementary Fig. 2a–d). We also examined the raw signals of BACS, which revealed a strong correlation between two biological replicates (Pearson’s *r* = 1.00, Supplementary Fig. 2e). Compared with the reported SILNAS mass spectrometry (SILNAS MS) data, 103 of 105 known Ψ sites in human cy-rRNA (including one 2’-*O*-methylpseudouridine, Ψm site in 28S rRNA) were identified with high confidence^44^ (Fig. 2c–e and Supplementary Fig. 2a–d). However, Ψ_1136_ in 18S rRNA was not detected, possibly due to its low modification level in HeLa cells (4.5% by BACS, Supplementary Fig. 2a,b). Interestingly, we found a 16% U-to-C mutation rate of the known 18S rRNA Ψ_36_ site in control libraries, although the mutation rate increased to 80% after BACS treatment (Fig. 2d and Supplementary Fig. 2f). Similar results were obtained from BID-seq control libraries^29^, suggesting the presence of an uncharacterized single nucleotide polymorphism (SNP) site in HeLa cells. It is noteworthy that these two sites could be readily detected in C666-1 cell line (Supplementary Fig. 2g). Therefore, BACS could detect all the known Ψ sites in human cy-rRNAs (Fig. 2f). In addition, we detected a new Ψ_4938_ site in 28S rRNA from HeLa and C666-1 cells, located adjacent to the previously known Ψ_4937_ site (Supplementary Fig. 2g). The presence of Ψ_4938_ was supported by two public databases, both of which predicted that small nucleolar RNA (snoRNA) SNORA17B would be responsible for catalyzing this modification^45,46^. BACS therefore provided further confirmation of the existence of the Ψ_4938_ site. We further identified four novel Ψ sites (Ψ_31_ / Ψ_890_ / Ψ_899_ in 18S rRNA and Ψ_1674_ in 28S rRNA) in cy-rRNAs from C666-1 cells (Supplementary Fig. 2g). It is important to note that while some Ψ or uridine modifications can induce intrinsic mutation signals (such as U-to-C mutation for m^1^acp^3^Ψ_1248_ in 18S rRNA and U-to-A mutation for m^3^U_4500_ in 28S rRNA)^47^, these can be filtered out by comparing results of BACS libraries with control libraries (Supplementary Fig. 2f). In addition to cy-rRNA, we also applied BACS to mitochondrial rRNA (mt-rRNA) and detected 8 and 1 Ψ sites in 12S and 16S rRNAs, respectively (Fig. 2b). Among them, 4 sites have also been detected by Pseudo-seq^4^. In general, the modification level of Ψ sites in mt-rRNA was significantly lower than their cytosolic counterparts (Supplementary Fig. 2h).

**Fig. 2.**
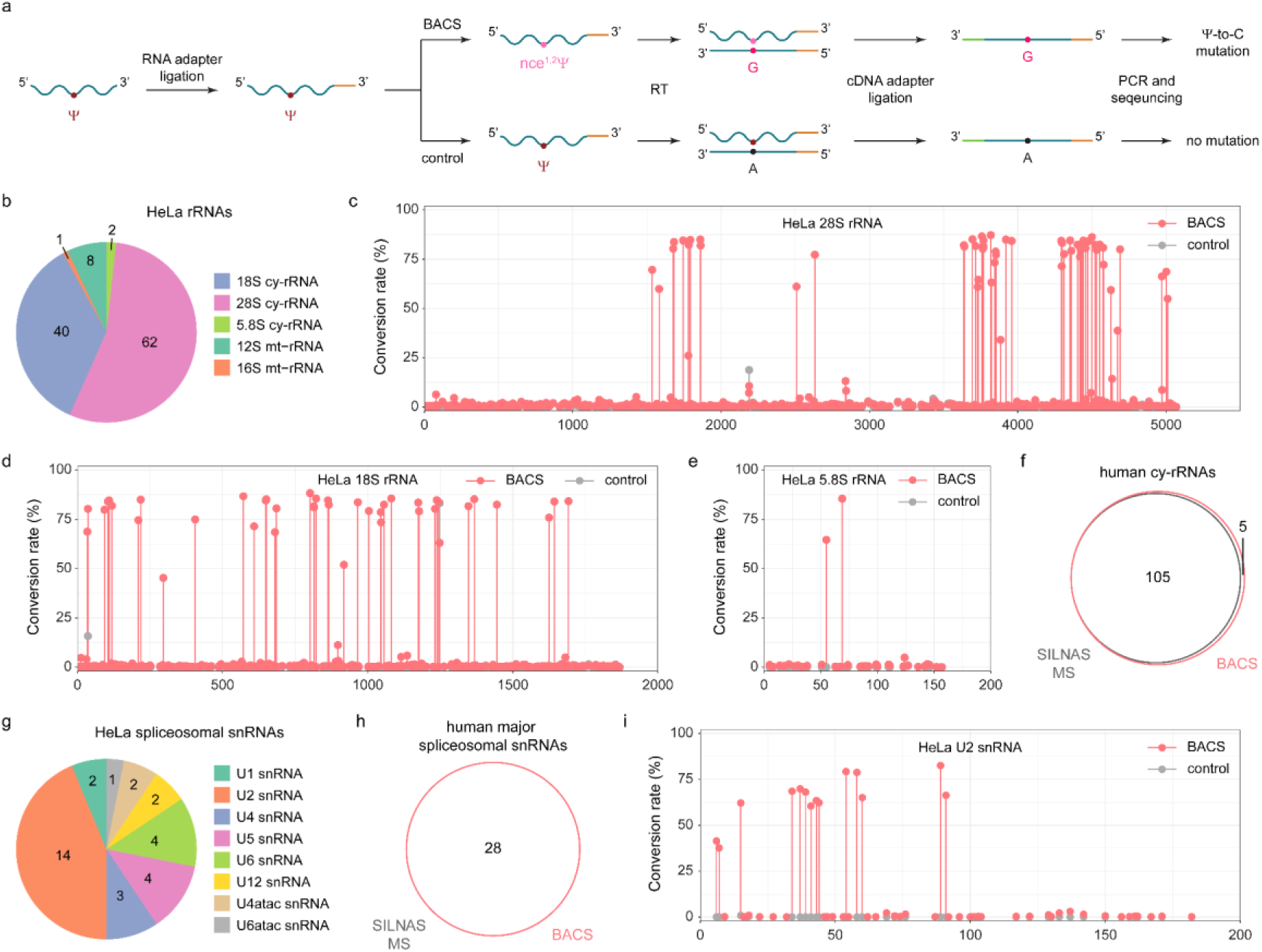
BACS detected known Ψ sites in human rRNA and spliceosomal snRNAs. **a.** Flowchart of BACS library construction. **b.** Numbers of Ψ sites identified in HeLa cy-rRNAs and mt-rRNAs. **c–e.** Conversion rates of BACS (pink) and control (grey) samples in HeLa 28S rRNA **(c)**, 18S rRNA **(d)**, and 5.8S rRNA **(e)**, respectively. Data are presented as means of two independent experiments. **f.** Venn diagram illustrating the overlap of Ψ sites detected in human cy-rRNAs between BACS and SILNAS MS. **g.** Numbers of Ψ sites identified in HeLa spliceosomal snRNAs. **h.** Venn diagram illustrating the overlap of Ψ sites detected in human spliceosomal snRNAs between BACS and SILNAS MS. **i.** Conversion rates of BACS (pink) and control (grey) samples in HeLa U2 snRNA. Data are presented as means of two independent experiments.

As expected, BACS clearly outperformed BS-based methods in the following aspects^29,30^. Firstly, the U-to-C mutation signature enabled BACS to determine the exact position and number of Ψ sites in consecutive uridine sequences (adjacent to one or more uridines, for instance, Ψ_801_ / Ψ_814_ / Ψ_815_ in 18S rRNA, Ψ_1847_ / Ψ_1849_ in 28S rRNA, and Ψ_4323_ / Ψ_4331_ in 28S rRNA), which remains challenging for BS-based methods (Supplementary Fig. 3a–c). The improved bioinformatics pipeline of BID-seq with realignment analysis (BID-pipe^31^) could not resolve ambiguity in two consecutive uridine sequences and may introduce artefacts in some cases (for example, Ψ_681_ in 18S rRNA, Ψ_1045_ / Ψ_1046_ in 18S rRNA, and Ψ_4549_ in 28S rRNA) (Supplementary Fig. 3d–f). These results demonstrate that the deletion signals induced by BS-based methods lack precision in consecutive uridine sequences and these issues are difficult to resolve by subsequent analysis. Secondly, more even conversion rates of Ψ sites were obtained over densely modified regions of rRNA using BACS compared with BS-based methods, indicating that BACS datasets would not be influenced by the density of pseudouridylation (Supplementary Fig. 3g,h). In particular, BACS successfully detected extremely dense Ψ sites in a narrow region (for example, Ψ_3737_ / Ψ_3741_ / Ψ_3743_ / Ψ_3747_ / Ψ_3749_ in 28S rRNA and Ψ_4263_ / Ψ_4266_ / Ψ_4269_ in 28S rRNA). Thirdly, when compared to SILNAS MS, BACS provided greatly improved accuracy in quantifying Ψ modification levels than BS-based methods (*r* = 0.90 for BACS vs *r* = 0.37 for BID-seq and *r* = 0.48 for PRAISE, Supplementary Fig. 3i–k). These findings strongly indicate that quantifying Ψ using deletion signals, as done by BS-based methods, introduces inaccuracies into the analysis. Finally, we found that BACS could achieve a higher conversion rate on 28S rRNA Ψm_3797_ site (85%) than BS-based methods (10– 20%), because BACS solely relied on the availability of *N*^1^ atom of Ψ (Supplementary Fig. 3h).

### BACS identified highly conserved Ψ sites in human spliceosomal snRNAs

After validating BACS on rRNA, we applied it to spliceosomal snRNAs from HeLa, C666-1, Raji, and Elijah cells, which contain multiple consecutive Ψ sites^48^. We first focused on major spliceosomal snRNA species and detected 2, 14, 3, 4, and 4 Ψ sites in U1, U2, U4, U5, and U6 snRNAs from HeLa cells, respectively, which is highly consistent with the latest SILNAS MS results^49^ (Fig. 2g). Only Ψ_59_ in U4 snRNA was not detected by BACS, since it is likely to be lowly modified in HeLa cells. However, this position was modified to higher levels in U4 snRNA from C666-1 and Elijah cells (Supplementary Fig. 4a). Consequently, we can detect all known Ψ sites in human major spliceosomal snRNAs (Fig. 2h). It is noteworthy that BACS successfully mapped all 14 Ψ sites in human U2 snRNA, which has not been realized by any other high-throughput sequencing methods, further highlighting the superiority of BACS in detecting dense and consecutive Ψ sites (Fig. 2i). Unlike the snRNA components of human major spliceosome, the Ψ profile of minor spliceosomal snRNA species has only been revealed using CMC-based primer extension assay, mainly due to their low abundance^50,51^. However, given that CMC-based methods may suffer from partial labeling efficiency and ‘stuttering’ phenomenon^15^, we believed that BACS could be a preferred method to study pseudouridylation in these snRNA species. Indeed, we consistently detected 2, 2, and 1 Ψ sites in U12, U4atac, and U6atac snRNAs from HeLa cells, respectively, while no Ψ site was detected in U11 snRNA (Fig. 2g). Notably, we confirmed that there are two consecutive Ψ sites (Ψ_11_ / Ψ_12_) rather than one Ψ_12_ site in U4atac snRNA, providing new insights into its interaction with U6atac snRNA^50,52^ (Supplementary Fig. 4b,c). Additionally, we detected highly conserved Ψ_247_ and Ψ_250_ sites in 7SK RNA^53^ and differentially modified Ψ_211_ site in 7SL RNA^49^ (Supplementary Fig. 4a). Our analysis also showed that there were no high-confidence Ψ sites in U7 snRNA, RNase P RNA, RNase MRP RNA, vault RNA, and Y RNA.

### BACS revealed the Ψ profile of human snoRNA

The Ψ profile of yeast snoRNA has been revealed through Pseudo-seq^4^ and Ψ-seq^5^, yet it remains relatively unexplored in human snoRNA. Ψ-seq^5^ and BID-seq^29^ only detected 11 and 39 Ψ sites in human snoRNA, respectively. In contrast, using BACS, we detected 304 Ψ sites in snoRNA from HeLa cells (Supplementary Fig. 5a). Further analysis revealed the presence of 205, 67, and 32 Ψ sites in box C/D snoRNAs, box H/ACA snoRNAs, and small Cajal body-specific RNAs (scaRNAs), respectively. Remarkably, all three types of snoRNAs exhibited a substantial number of highly modified Ψ sites (Supplementary Fig. 5b,c). SnoRNA Ψ sites detected through BACS largely covered those previously identified by Ψ-seq^5^ and BID-seq^29^, demonstrating increased sensitivity of BACS (Supplementary Fig. 5d,e). Furthermore, we observed that Ψ sites in box C/D snoRNAs displayed enrichment in the 5’-upstream regions of box D’ and the 3’-downstream regions of box C’, while Ψ sites in box H/ACA snoRNAs were enriched in the 5’-upstream regions of box H and ACA (Supplementary Fig. 6a,b). These patterns implied a potential role for Ψ in mediating interactions between snoRNAs and their targets. Indeed, a subset of Ψ sites identified in box C/D and box H/ACA snoRNAs were also located in the predicted guide regions, which was in accordance with the Ψ-seq results^5^ (Supplementary Fig. 6c,d).

In addition, human telomerase RNA component (TERC) shares similar characteristics with snoRNAs, as it contains a conserved box H/ACA scaRNA domain at the 3’-end^54^. Upon BACS treatment, we detected 7 Ψ sites in TERC from HeLa cells, 4 of which were putative Ψ sites previously discovered through CMC-based primer extension approach^55^ (Supplementary Fig. 5a,f). In particular, all 3 novel Ψ sites (Ψ_38_ / Ψ_100_ / Ψ_155_), together with the known Ψ_161_ and Ψ_179_ sites, were found in the core domain of TERC. This observation suggested the potential involvement of Ψ in stabilizing the TERC structure, similar to the scenario that has been demonstrated for Ψ_306_ and Ψ_307_ within the P6.1 loop of TERC^56^.

### A comprehensive Ψ map of human tRNA

Ψ is one of the most fundamental and prevalent modifications in human tRNA^57,58^. However, given that most of tRNA species are extensively modified and highly structured, quantitative profiling of Ψ in tRNA remains challenging by CMC- or BS-based methods^30,59^. We anticipated that BACS would offer a better solution to this problem, since mutation signals induced by BACS would not be influenced by RT blocks or other intrinsic mismatches. Indeed, while CMC- and BS-based methods failed to map human cytosolic tRNAs (cy-tRNAs), we used BACS to generate the first quantitative Ψ map of cy-tRNAs from HeLa cells with 609 high-confidence Ψ sites (Supplementary Fig. 7a). The number of Ψ sites identified per cy-tRNA varied among different isotypes (Fig. 3a). In cy-tRNAs, Ψ sites were predominantly located at highly conserved positions, including position 13, 27–28, 38–40, and 55, while Ψ at other positions were limited to specific types of cy-tRNAs (Fig. 3b). An integrated view of the Ψ profile of human cy-tRNAs was then summarized based on the canonical tRNA numbering system^60^ (Supplementary Fig. 7b). Subsequently, we compared the Ψ modification level at each tRNA position (Fig. 3c). Notably, position 55 emerged as the most frequently and highly modified Ψ site in cy-tRNAs. Moreover, position 13 also displayed a high level of Ψ modification. In contrast, the modification levels of position 27–28 and position 38–40 showed considerable variations.

**Fig. 3.**
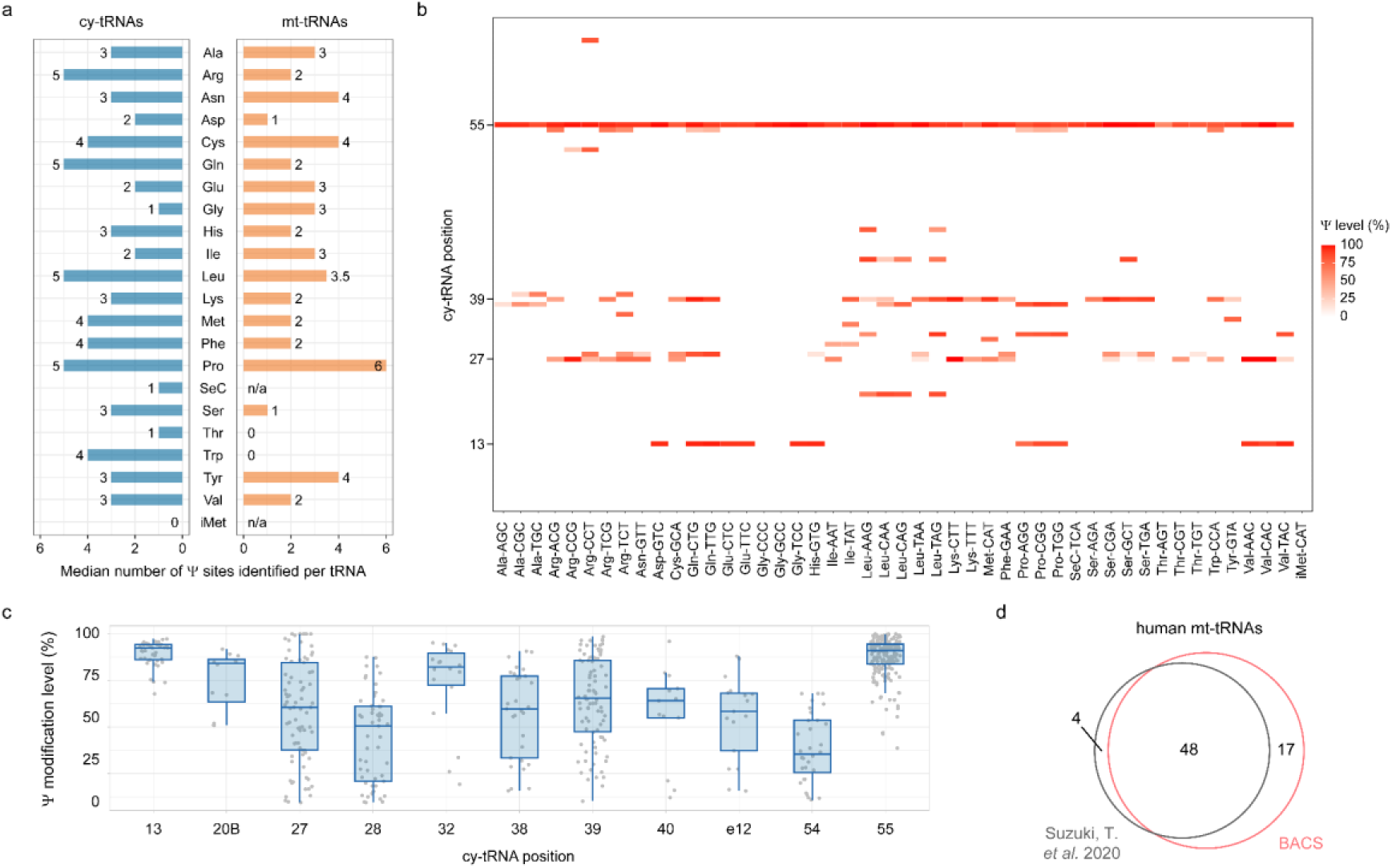
BACS unveiled the comprehensive Ψ profile of human tRNA. **a.** Median numbers of Ψ sites identified per tRNA in each cy-tRNA (left) and mt-tRNA (right) isotype from HeLa cells. n/a, not applicable. **b.** Heatmap showing the modification levels of high-confidence Ψ sites in HeLa cy-tRNAs. Only one representative tRNA isodecoder was presented for each isoacceptor family. **c.** Comparison of the modification levels of Ψ sites at selected positions of HeLa cy-tRNAs. Boxplots visualize all Ψ sites at each position, indicating medians, quantiles, and extreme values (tRNA position: 13, *n* = 43; 20B, *n* = 12; 27, *n* = 81; 28, *n* = 53; 32, *n* = 18; 38, *n* = 29; 39, *n* = 86; 40, *n* = 13; e12, *n* = 17; 54, *n* = 32; 55, *n* = 185). **d.** Venn diagram illustrating the overlap of Ψ sites in human mt-RNAs reported by BACS and a previously published dataset^61^.

Using BACS, we also detected 54 Ψ sites in HeLa mitochondrial tRNAs (mt-tRNAs) (Fig. 3a and Supplementary Fig. 7a). Applying BACS to other human cell lines, we observed significant differential Ψ modification on mt-tRNAs. For example, Ψ_55_ in mt-tRNA^Met^ was not characterized as high-confidence sites due to its low modification level in HeLa cells (3.6%), while it was readily detected in C666-1, Raji, and Elijah cell lines (Supplementary Fig. 7c,d). In addition, C666-1 cell lines displayed an elevated Ψ level at position 66–68 compared to HeLa, Raji, and Elijah cells (Supplementary Fig. 7d). We finally mapped a total of 65 Ψ sites in mt-tRNAs by merging the results of HeLa, C666-1, Raji, and Elijah cells, which was highly consistent with the published dataset^61^ (Fig. 3d). Overall, human mt-tRNAs were pseudouridylated to a lesser extent compared with cy-tRNAs (Fig. 3b,c and Supplementary Fig. 7c,d). In contrast, PRAISE detected only 34 Ψ sites in mt-tRNAs from HEK293T cells (Supplementary Fig. 8a). PRAISE encountered challenges in quantifying consecutive Ψ sites (8 out of 34, 23.5%) and determining the precise position for single Ψ site within multiple uridine contexts (13 out of 34, 38.2%) (Supplementary Fig. 8b). The signal produced by PRAISE for these Ψ sites manifested as an ambiguous peak, lacking accurate quantification and single-base resolution (Supplementary Fig. 8c–e). As a result, PRAISE achieved quantitative and single-base resolution detection for only 13 Ψ sites in mt-tRNAs, revealing significant limitations compared to BACS.

### Profiling and quantification of Ψ in HeLa mRNA

After successfully applying BACS to various types of ncRNAs, we extended its usage to map and quantify Ψ modifications in HeLa mRNA. Given that the low stoichiometry of Ψ modification in polyA-tailed RNA, we applied in vitro transcribed polyA-tailed RNA (IVT RNA) from HeLa cells as a modification-free control to help with Ψ calling^62^ (Supplementary Fig. 9a). We detected a total of 1335 Ψ sites in HeLa polyA-tailed RNA (Fig. 4a). The majority of these Ψ sites exhibited low modification levels (<20%), while only a limited number of Ψ sites displayed high levels of modification (>50%) (Fig. 4a,b). In contrast to the aforementioned ncRNAs, the Ψ modification level in polyA-tailed RNA was significantly lower (Supplementary Fig. 9b). Among the 1335 Ψ sites, 1294 and 41 were located in mRNA and ncRNA (excluding rRNA, snRNA, snoRNA, and tRNA), respectively (Fig. 4c). Within mRNA, Ψ was enriched in the coding sequence (CDS) and 3’-untranslated region (3’-UTR), while it was relatively depleted in the 5’-untranslated region (5’-UTR), consistent with previous findings^20,29,30^ (Fig. 4c,d). The 1335 Ψ sites were located across 1103 polyA-tailed RNA transcripts, with the majority carrying only one Ψ site (Fig. 4e). Gene ontology (GO) analysis revealed that Ψ-modified mRNA was enriched in functions such as translation and regulation of apoptotic process (Supplementary Fig. 9c). Importantly, BACS could simultaneously provide mRNA expression levels while mapping Ψ, which showed strong correlation with control libraries (Pearson’s *r* = 0.99– 1.00), suggesting minimal RNA degradation induced by BACS (Supplementary Fig. 9d).

**Fig. 4.**
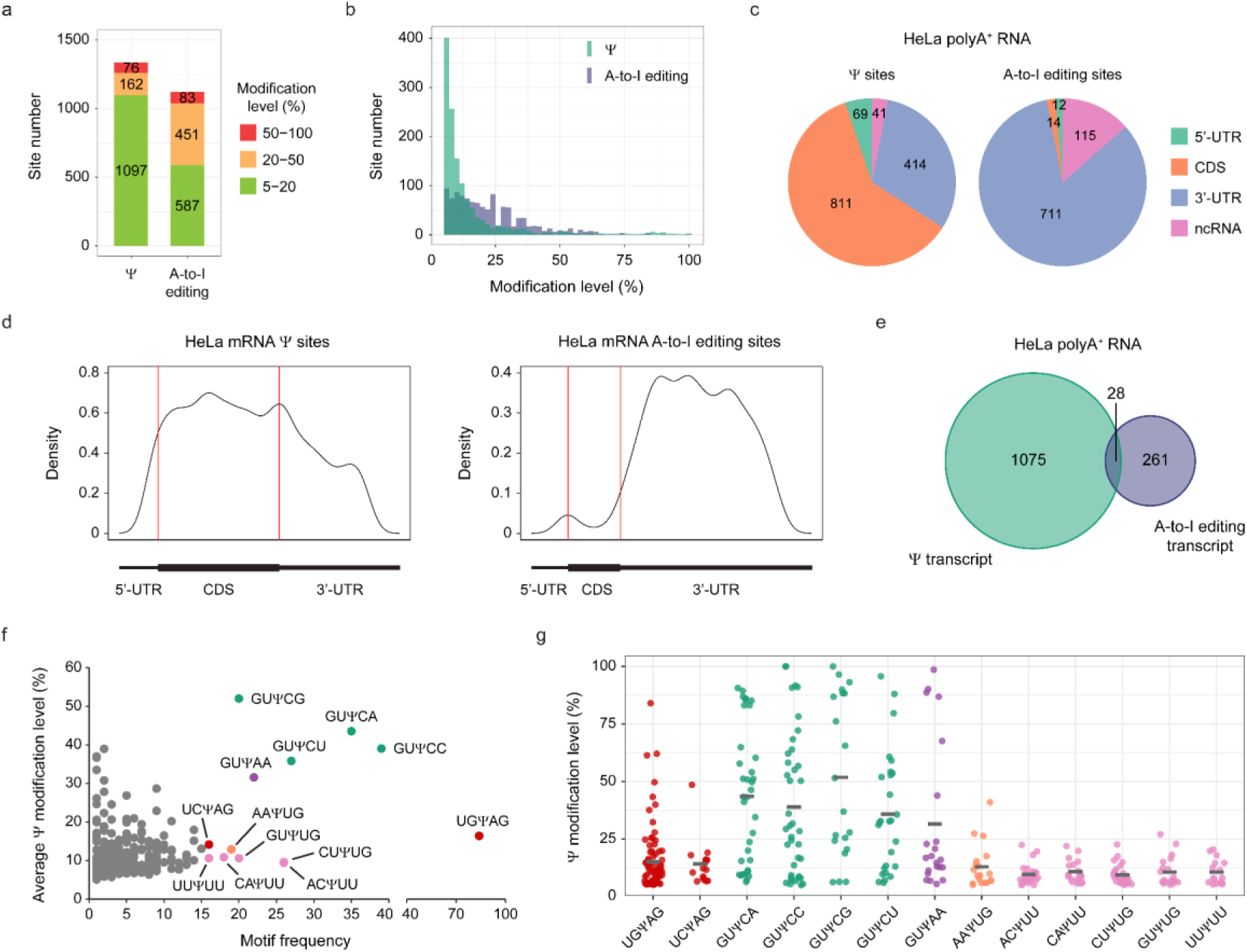
Simultaneous characterization of Ψ and A-to-I editing sites in HeLa mRNA. **a.** Numbers of Ψ and A-to-I editing sites with high (50–100%, red), medium (20–50%, yellow), and low (5–20%, green) modification levels identified in HeLa polyA-tailed RNA. **b.** Modification level distribution of Ψ and A-to-I editing sites in HeLa polyA-tailed RNA. **c.** Distribution of Ψ and A-to-I editing sites within different features of HeLa mRNA and ncRNA. **d.** Metagene profile of Ψ and A-to-I editing sites in HeLa mRNA. **e.** Venn diagram illustrating the overlap of transcripts possessing Ψ and A-to-I editing sites. **f.** Motif frequency of Ψ sites in HeLa mRNA. **g.** Modification level distributions of HeLa mRNA Ψ sites within selected motifs, with means indicated in each plot by a horizontal line. Motif: UGΨAG, *n* = 84; UCΨAG, *n* = 16; GUΨCA, *n* = 35; GUΨCC, *n* = 39; GUΨCG, *n* = 20; GUΨCU, *n* = 27; GUΨAA, *n* = 22; AAΨUG, *n* = 19; ACΨUU, *n* = 26; CAΨUU, *n* = 18; CUΨUG, *n* = 26; GUΨUG, *n* = 20; UUΨUU, *n* = 16.

Next, we analyzed the sequence contexts of Ψ in HeLa mRNA. First, our analysis indicated that the majority of Ψ sites (55.3%) were located in consecutive uridine sequences (Supplementary Fig. 10a). These positions could not be precisely determined through BS-based methods^29,30^, further highlighting the advantage of BACS. Benefiting from the high-resolution signals of BACS, we found that Ψ was predominantly enriched in USΨAG (S = C or G) and GUΨCN (N = A, C, G or U) motifs, corresponding to the previously identified PUS7 and TRUB1 motif, respectively^63^ (Fig. 4f). In addition, we also observed that Ψ tends to be enriched in those motifs containing multiple consecutive uridines, such as CUΨUG, ACΨUU, and even UUΨUU. The stoichiometry of Ψ within these motifs was also compared, demonstrating that GUΨCN exhibited a relatively high modification level (Fig. 4g). Furthermore, we analyzed the codon preference of Ψ in mRNA. As expected, Ψ was enriched in those codons containing consecutive uridines, such as UUY (Y = C or U), UUG, AUU, and GUU, which encoded phenylalanine (Phe), leucine (Leu), isoleucine (Ile), and valine (Val), respectively (Supplementary Fig. 10b,c). Within codons, Ψ was mainly located in the second position (Supplementary Fig. 10d). Moreover, we observed one Ψ site positioned in the start codon (AUG), while two sites were found in the stop codon (UAG), which may promote stop codon readthrough according to previous research^29,64^.

To further evaluate the performance of BACS, we compared our identified mRNA Ψ sites with published datasets. We first compared BACS with a recent dataset which consolidated three CMC-based methods^63^. Remarkably, BACS accurately identified 61 out of 70 Ψ sites (87.1%) listed in the “highest confidence” category (Supplementary Fig. 11a). However, a strong overlap between BACS and the “high confidence” list was achieved only when considering Ψ sites consistently detected across multiple samples (>8) (177 out of 320 Ψ sites, 55.3%), indicating the considerable variance between different CMC-based datasets (Supplementary Fig. 11b,c). We further compared BACS with two recently developed BS-based methods. Compared to CMC-based approaches, BACS demonstrated a better overlap with BID-seq results, as expected^29^ (230 out of 575 Ψ sites, 40.0%, Supplementary Fig. 11d). Most of the sites exclusive to the BID-seq dataset displayed low modification levels in our BACS libraries, possibly due to the inaccurate quantification of BID-seq (Supplementary Fig. 11e). When compared with PRAISE^30^, 651 of 1995 Ψ sites (32.6%) showed an overlap with BACS results (Supplementary Fig. 11f). Similarly, the majority of PRAISE-only Ψ sites were lowly modified in our dataset (Supplementary Fig. 11g). Regarding the seven Ψ sites identified in mitochondrial mRNAs (mt-mRNAs) by BACS, four, two, and four of them have also been detected by Pseudo-seq^4^, BID-seq^29^, and PRAISE^30^, respectively. It is important to note that the degree of overlap between different methods may be influenced by differences in sequencing depth and the bioinformatics pipelines used for analysis (for example, mapping to the genome or directly to the transcriptome). Therefore, there is a clear need for developing a standardized pipeline for analyzing RNA modifications.

### Simultaneous profiling of A-to-I editing sites and m^1^A in HeLa RNA

In addition to Ψ mapping, BACS enabled the simultaneous detection of A-to-I editing sites. After BACS, the A-to-G mutation signals induced by A-to-I editing would be erased in a similar way as shown in ICE-seq, which could distinguish the true editing sites from potential backgrounds^65^ (Supplementary Fig. 9a). We identified 1121 A-to-I editing sites in HeLa polyA-tailed RNA, with a mean modification level of 20% (Fig. 4a,b). In stark contrast to Ψ, the majority of A-to-I editing sites were resident in the *Alu* elements (Supplementary Fig. 9e). We further annotated 737 and 115 A-to-I editing sites to mRNA and ncRNAs, respectively (Fig. 4c). Within mRNA, the A-to-I editing sites were predominantly enriched in 3’-UTR, consistent with previous findings (Fig. 4d)^66^. Interestingly, mRNA transcripts that carry A-to-I editing sites did not overlap with those possessing Ψ, suggesting distinct roles of A-to-I editing and pseudouridylation in mRNA processing (Fig. 4e).

Similar to RBS-seq^25^, BACS induces Dimroth rearrangement of m^1^A to *N*^6^-methyladenosine (m^6^A) and could potentially detect m^1^A together with Ψ (ref.^67^) (Supplementary Fig. 12a). As expected, we observed a significant reduction of m^1^A mutation signals at tRNA position 58 (for cy-tRNAs) and 9 (for mt-tRNAs) after BACS treatment, which was comparable to the efficiency of demethylase^68^ (Supplementary Fig. 12b,c). These results highlight the value of BACS to detect multiple modifications simultaneously.

### BACS uncovered novel PUS targets and activities in HeLa cells

To elucidate the PUS-dependent Ψ profile in the HeLa transcriptome, we generated individual knock out for three key PUS enzymes: TRUB1, PUS7, and PUS1 (Fig. 5a). In contrast to the conventional view that TRUB1 was the sole PUS enzyme responsible for cy-tRNA Ψ55 (ref.^69^), depletion of TRUB1 did not eradicate Ψ55 in human cy-tRNAs, suggesting that other PUS enzymes (likely TRUB2 and PUS10) may also participate in the pseudouridylation of this position (Fig. 5b,c). Surprisingly, all the Ψ55 sites in mt-tRNA were eliminated upon TRUB1 depletion, challenging another conventional belief that TRUB2 was solely responsible for mt-tRNA Ψ55 (ref.^70^) (Fig. 5b,c). We further discovered that the TRUB1 motif would extend beyond the recognized GUΨCNA (N = A, C, G or U) motif^5,63^, since it could modify GUΨUAA in mt-tRNA^Asn^, GUΨGUA in mt-tRNA^Glu^, and GUΨAAA in mt-tRNA^Pro^ with high efficiency (Fig. 5d and Supplementary Fig. 13a–c). We observed similar results in polyA-tailed RNA, confirming that TRUB1 could edit GUΨGNA, GUΨCNA and GUΨUNA motifs (Fig. 5d). Therefore, we demonstrate that the TRUB1 motif can be extended from GUΨCNA to GUΨNNA.

**Fig. 5.**
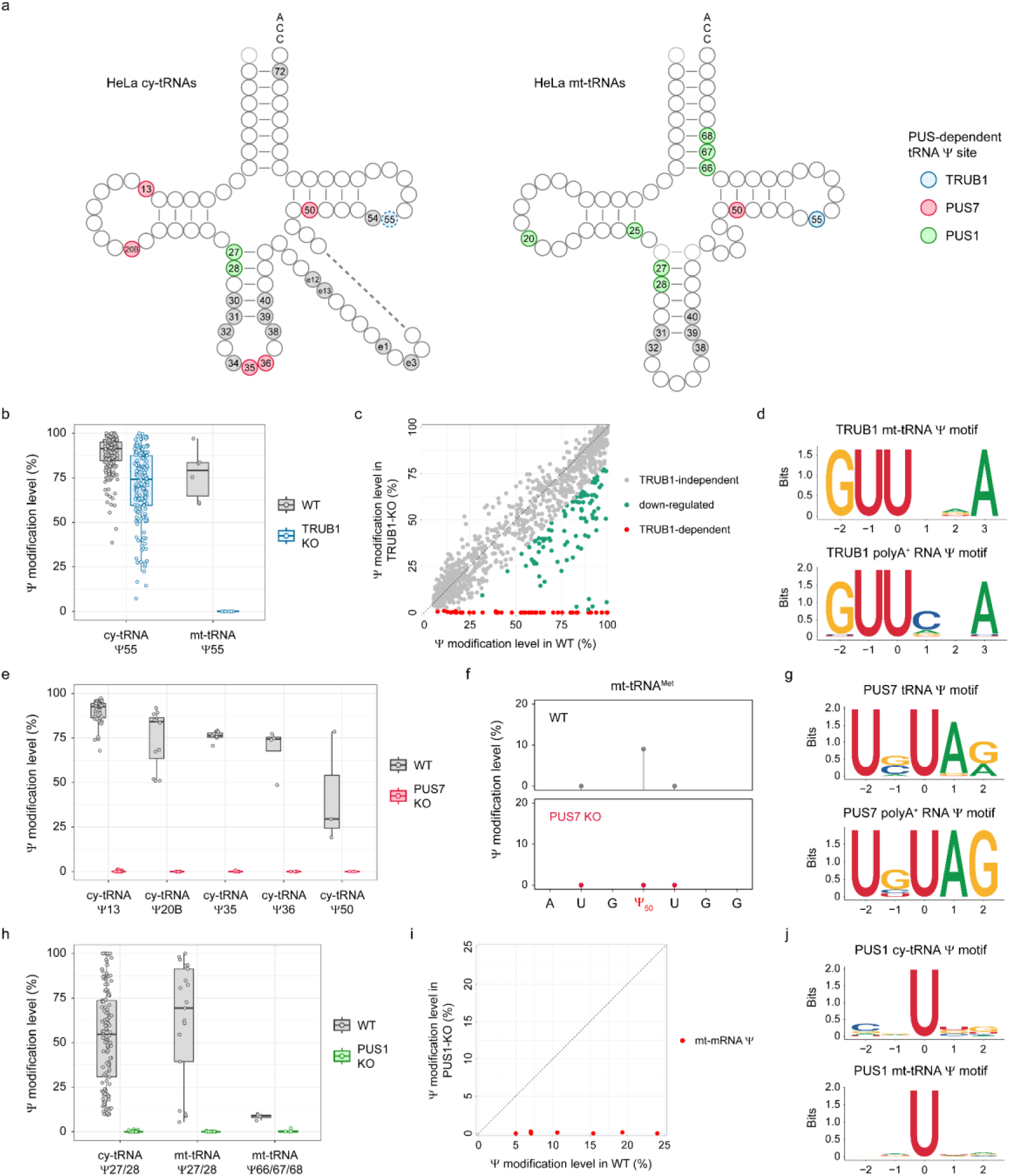
PUS-dependent Ψ landscape across the HeLa transcriptome. **a.** Integrated view of the PUS-dependent Ψ profiles of HeLa cy-tRNAs (left) and mt-tRNAs (right). Blue, red, and green color denote the Ψ targets of TRUB1, PUS7, and PUS1, respectively. Ψ55 in HeLa cy-tRNAs is partially dependent on TRUB1, which is labeled by dashed line. **b.** Comparison of the modification levels of Ψ55 in HeLa cy-tRNAs and mt-tRNAs upon TRUB1 depletion. Boxplots visualize all Ψ sites at each position, indicating medians, quantiles, and extreme values (cy-tRNA Ψ55, *n* = 180; mt-tRNA Ψ55, *n* = 6). **c.** Scatter plot illustrating all TRUB1-dependant Ψ sites across the HeLa transcriptome. **d.** Sequence motifs of TRUB1-dependent Ψ sites in mt-tRNAs (upper panel) and polyA-tailed RNA (lower panel). **e.** Comparison of the modification levels of Ψ sites at selected positions of HeLa cy-tRNAs upon PUS7 depletion. Boxplots visualize all Ψ sites at each position, indicating medians, quantiles, and extreme values (cy-tRNA: Ψ13, *n* = 43; Ψ20B, *n* = 12; Ψ35, *n* = 7; Ψ36, *n* = 4; Ψ50, *n* = 3). **f.** Comparison of the modification levels of Ψ50 in mt-tRNA^Met^ between WT and PUS7-KO cell lines. **g.** Sequence motifs of PUS7-dependent Ψ sites in tRNAs (upper panel) and polyA-tailed RNA (lower panel). **h.** Comparison of the modification levels of Ψ sites at selected positions of HeLa cy-tRNAs and mt-tRNAs upon PUS1 depletion. Boxplots visualize all Ψ sites at each position, indicating medians, quantiles, and extreme values (cy-tRNA: Ψ27/28, *n* = 130; mt-tRNA: Ψ27/28, *n* = 21; Ψ66/67/68, *n* = 4). **i.** Scatter plot illustrating all PUS1-dependant Ψ sites in HeLa mt-mRNAs. **j.** Sequence motifs of PUS1-dependent Ψ sites in cy-tRNAs (upper panel) and mt-tRNAs (lower panel).

PUS7-knockout HeLa cells allowed us to reveal its role in the formation of Ψ20B, Ψ36, and Ψ50 in cy-tRNAs, in addition to its previously known targets Ψ13 and Ψ35 (ref.^71^) (Fig. 5e and Supplementary Fig. 13d). It is noted that both Ψ35 and Ψ36 are located within the anticodon, suggesting that PUS7 depletion could induce miscoding events of cy-tRNA^Tyr(GTA)^ and cy-tRNA^Arg(TCT)^, respectively. Moreover, our study uncovers a novel activity for PUS7 in mitochondria by catalyzing the modification of Ψ50 in mt-tRNA^Met^, expanding its functional repertoire within this organelle for the first time (Fig. 5f). The majority of PUS7 targets in tRNA displayed conserved UVΨAR (V = A, C or G; R = A or G) motif (Fig. 5g). However, PUS7 displayed comparable activity within the UGΨGG motif (cy-tRNA^Arg(CTT)^ Ψ50) and relatively low activity within the UGΨUG motif (mt-tRNA^Met^ Ψ50) (Fig. 5f,g and Supplementary Fig. 13e). In polyA-tailed RNA, PUS7 mainly catalyzed the pseudouridylation within UBΨAG (B = C, G or U) motif (Fig. 5g). Collectively, the true PUS7 consensus motif would be UNΨAR (N = A, C, G or U; R = A or G) and UGΨKG (K = G or U), which was less strict than previously considered UGΨAR (R = A or G) motif^4,72^.

Finally, PUS1 depletion resulted in the complete loss of Ψ27/28 in human cy-tRNAs (Fig. 5h and Supplementary Fig. 13f). In mt-tRNAs, PUS1 not only catalyzed the modification of Ψ27/28 but also induced the formation of Ψ66/67/68, consistent with the findings on yeast and mouse PUS1 homologs^73^ (Fig. 5h and Supplementary Fig. 13f). We also observed non-canonical activity of PUS1, exemplified by Ψ25 in mt-tRNA^Asn^ and Ψ20 in mt-tRNA^Leu(UUR)^, thus affirming the precision of BACS in capturing diverse pseudouridylation events (Supplementary Fig. 13g,h). Additionally, PUS1 was found to catalyze all 7 Ψ sites identified in mt-mRNAs (Fig. 5i). These results show that PUS1 is the major PUS enzyme in mitochondria with diverse functions. PUS1-dependent Ψ sites did not show any sequence motifs, consistent with an earlier report that its activity is dependent on RNA structure^72^ (Fig. 5j).

Mimicry of tRNA has been considered as a general way for mRNA pseudouridylation^5,63^. Although TRUB1, PUS7, and PUS1 could edit both tRNAs and polyA-tailed RNA, the Ψ targets in polyA-tailed RNA were significantly less modified than their counterparts in tRNAs (Supplementary Fig. 13i). These results suggest that mRNA may not be the primary substrate of these stand-alone PUS enzymes, which is consistent with our earlier data showing that the majority of mRNA Ψ sites were modified to a low level.

### Mapping of Ψ in viral RNAs

It has been widely accepted that Ψ-modified RNAs can suppress innate immune responses and may influence the mRNA vaccine design^74^. Although several studies have reported the presence of Ψ in viral RNAs^75,76^, it has not been thoroughly confirmed with the latest sequencing technologies. We therefore applied BACS to various RNA viruses, including severe acute respiratory syndrome coronavirus 2 (SARS-CoV-2), hepatitis C virus (HCV), Zika virus (ZIKV), hepatitis delta virus (HDV), and Sindbis virus (SINV), to study the existence of Ψ in viral transcripts and genomes. We first focused on SARS-CoV-2, since earlier studies reported Ψ in its genomic and subgenomic RNAs by Nanopore sequencing^75,76^. Similar to previous RNA-seq results^77^, the majority of reads were mapped to the positive strand with a unique pattern at the 3’-end corresponding to the subgenomic RNAs (Supplementary Fig. 14a). However, we could not detect any high-confidence Ψ sites in SARS-CoV-2 RNA (Fig. 6a and Supplementary Fig. 14b). Surprisingly, we did not identify any high-confidence Ψ sites in the other four RNA viruses (Supplementary Fig. 14c–f). Importantly, we confirmed robust viral infection in our model systems, to ensure a high abundance of viral RNAs concomitant with high depth of coverages (Supplementary Table 3). These results suggest that Ψ is not directly involved in the modification of these RNA viruses. In addition to these RNA viruses, we also applied BACS to human cell lines infected by Epstein-Barr virus (EBV), a DNA virus that encodes two highly expressed ncRNAs, EBER1 and EBER2, both of which are RNA polymerase III transcripts^78,79^. A previous study using HydraPsi-seq^80^ and CMC-based primer extension methods reported one lowly modified Ψ_160_ site in EBER2 (ref.^81^) (Fig. 6b). However, we detected one highly modified Ψ_114_ site in EBER2, while no Ψ site was identified in EBER1, indicating the previous results were likely caused by high backgrounds from hydrazine and CMC chemistry (Fig. 6b,c). Notably, this novel Ψ site was predicted to be in a loop region accessible for PUS enzymes. The EBER2 Ψ_114_ site was conserved across all EBV strains and host cell lines tested, including nasopharyngeal carcinoma cell line C666-1 and two Burkitt’s lymphoma cell lines Raji and Elijah (Fig. 6c). Taken together, these results suggest different ways of utilizing Ψ between virus families and further highlight BACS as a highly specific method.

**Fig. 6.**
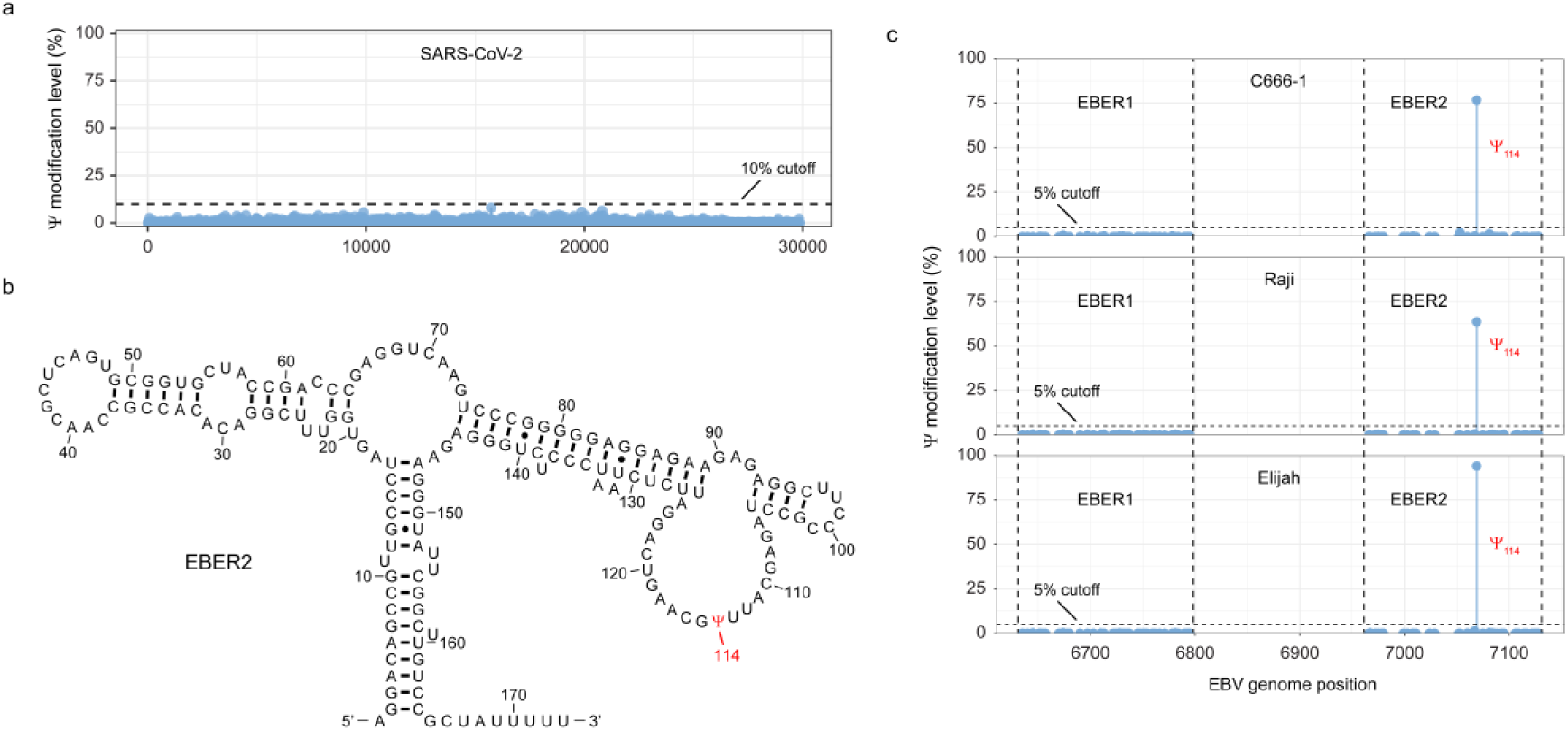
Investigation of Ψ modification in viral RNAs. **a.** Ψ modification levels in SARS-CoV-2 viral RNA. **b.** Canonical EBER2 structure, with Ψ_114_ site labeled accordingly. EBER2 structure is adapted from Rfam^108^. **c.** Ψ modification levels in EBER1 and EBER2 from C666-1 (upper panel), Raji (middle panel), and Elijah (lower panel) cell lines.

## Discussion

In this study, we report the development of a novel method, named BACS, for quantitative and base-resolution sequencing of Ψ. BACS is based on new bromoacrylamide cyclization chemistry and induces Ψ-to-C mutation signatures rather than truncation or deletion signatures, allowing for accurate quantification of Ψ stoichiometry and sequencing of Ψ at absolute single-base resolution. Importantly, BACS overcomes the inherent limitations of BS-based methods in three crucial aspects: (1) it facilitates the precise determination of Ψ sites located adjacent to one or more uridines; (2) it enhances the detection of densely modified Ψ sites with higher accuracy and sensitivity; and (3) it offers significantly more accurate quantification of Ψ in all sequence contexts. These advances make BACS a valuable tool for studying Ψ modifications in cellular RNAs, as it can provide a more comprehensive and accurate picture of the Ψ landscape across various RNA species. Using BACS, we successfully detected all known Ψ sites in human rRNA and spliceosomal snRNAs and generated the first quantitative Ψ map of human snoRNA and tRNA. Combining BACS with the latest library construction techniques has the potential to further improve its performance on small RNAs^68,82^. We further applied BACS to HeLa mRNA and revealed a rather low level of pseudouridylation. Moreover, by genetic depletion of PUS enzymes, we identified novel targets of TRUB1, PUS7, and PUS1 in HeLa cells, demonstrating their involvement in more diverse biological processes than previously thought. The absolute single-base resolution of BACS enabled us to extend the sequence motifs of TRUB1 and PUS7. Our results suggest at least one redundant PUS enzyme for TRUB1 editing of cy-tRNA Ψ55 modification, while no enzyme redundancy was found for PUS1 and PUS7. Given that most of previous studies on PUS enzymes were conducted on yeast^8^, our results indicate a broader range of activities for human PUS enzymes compared to their yeast homologues. There is a clear need to revisit the activities of human PUS enzymes on ncRNAs instead of relying solely on yeast results. BACS could serve as a valuable tool for studying PUS knockout cells to elucidate the properties and functions of the other 10 PUS enzymes. Finally, we mapped the Ψ landscape of EBV-encoded ncRNAs EBER1 and EBER2 and showed that several human RNA viruses (SRAS-CoV-2, HCV, ZIKV, HDV, and SINV) lack Ψ modifications in their transcripts or genomes.

Indeed, while significant progress has been made in understanding the function of Ψ, there are still many unknowns surrounding this abundant yet enigmatic modification^83^. The scarcity of Ψ in mRNA compared to ncRNAs has raised questions about its functional significance in mRNA. However, recent research has highlighted that Ψ could be much more abundant in pre-mRNA, suggesting that it could play important roles during pre-mRNA processing^84^. BACS could be well-suited to investigate the pseudouridylation of nascent RNA.

In light of these potential applications, we anticipate BACS to be widely adopted as the new standard to advance our understanding of Ψ modifications and their functional implications in diverse biological processes.

## Methods

### Preparation of model RNA

Regular and Ψ-labeled 10mer RNA oligonucleotides and 30mer spike-ins were purchased from Integrated DNA Technologies (IDT). The 72mer Ψ-containing model RNA used for mutation analysis and the 1.8-kb 10% Ψ-modified RNA used for UHPLC-MS/MS were prepared by T7 *in vitro* transcription using HiScribe T7 High Yield RNA Synthesis Kit (NEB) and Pseudo-UTP (Jena Bioscience) according to the manufacturer’s protocol. The DNA template was removed by adding 2 μl Turbo DNase (Invitrogen) to the reaction and incubating at 37 °C for 30 min. The products were finally purified with Monarch RNA Cleanup Kit (NEB). Sequences of RNA oligonucleotides can be found in Supplementary Table 1.

### Mass spectrometry analysis of short oligonucleotides

MALDI was performed on a Voyager-DE Biospectrometry Workstation (Applied Biosystems) with 2’,4’,6’-trihydroxyacetophenone (THAP) as matrix. All the oligonucleotides were analyzed in positive mode.

### Quantification of Ψ level by UHPLC-MS/MS

The untreated and treated RNA were digested into nucleosides by Nucleoside Digestion Mix (NEB) in a 50 µl solution according to the manufacturer’s protocol. After filtering with Amicon Ultra-0.5 mL 3K centrifugal filters (Millipore), the digested samples were subjected to UHPLC–MS/MS analysis as described before^85^. 1290 Infinity LC Systems (Agilent) was equipped with a ZORBAX RRHD SB-C18 column (2.1 × 150 mm, 1.8 μm, Agilent) coupled with a 6495B Triple Quadrupole Mass Spectrometer (Agilent). The ions were monitored in positive mode with mass transitions of m/z 245 to 125 (Ψ+H) and m/z 245 to 113 (rU+H) (Supplementary Table 2). Concentrations of nucleosides in RNA samples were deduced by fitting the signal peak areas into the standard curves.

### Cell culture

HeLa cells were cultured in DMEM medium (Gibco) supplemented with 10% (v/v) FBS (Gibco) and 1% penicillin/streptomycin (Gibco) at 37 °C with 5% CO_2_. For isolation of RNA, cells were harvested by centrifugation for 5 min at 1,000× g and room temperature.

C666-1, Raji, and Elijah cells were grown in Roswell Park Memorial Institute 1640 medium (RPMI 1640) (Thermo), complemented with 10% (v/v) fetal bovine serum (FBS) (Biosera), 2 mM L-glutamine (Thermo), and 100 units/ml penicillin and 100 µg/ml streptomycin (Thermo). Cells were grown in humidified incubator at 37 °C with 5% CO_2_. C666-1 cell line was gifted from Dr Christopher Dawson (University of Warwick). Raji cell line was gifted from Prof. Paul Farrell (Imperial College London). Cell lines were monthly tested for mycoplasma using MycoAlert Kit (Lonza) and were sent for authentication by Eurofins genomics.

### Generation of CRISPR knockout cell lines

TRUB1-KO, PUS7-KO, and PUS1-KO HeLa cells were generated using CRIScoPR-Cas9 technology (Supplementary Fig. 15). Briefly, single guide RNA (sgRNA) sequences were cloned into PX459 plasmids^86^. Transfection was performed using Lipofectamine 3000 Transfection Reagent (Invitrogen) following the manufacturer’s protocol. Cells were then selected by 2 µg/ml puromycin (Thermo). Serial dilution was performed to achieve clonal isolation. Finally, clones were expanded and picked for western blot validation with TRUB1 antibody (Proteintech, #12520-1-AP), PUS7 antibody (Abcam, #ab226257), and PUS1 antibody (Proteintech, #11512-1-AP), respectively. The sgRNA sequences were listed as follows:

TRUB1: 5’-CACGGCGAACACGCCGCTCAAGG-3’;
PUS7-sgRNA1: 5’-TTAATATTGAAACCCCGCTCTGG-3’;
PUS7-sgRNA2: 5’-TCGGAATGCAGTCTAACCAAAGG-3’;
PUS1: 5’-AATACAGCCTGACCGGACGAGGG-3’.

### RNA isolation

Total RNA was isolated using TRIzol (Invitrogen) and Direct-zol RNA Miniprep Plus (Zymo Research) according to the manufacturer’s protocol. Ribo^−^ RNA was isolated using RiboMinus Eukaryote System v2 (Invitrogen) according to the manufacturer’s protocol. PolyA^+^ RNA was isolated by two rounds of polyA-tailed selection using Dynabeads mRNA DIRECT Purification Kit (Invitrogen) according to the manufacturer’s protocol. To remove genomic DNA contamination, RNA was then treated with Turbo DNase and purified by Zymo-IC Column with RNA binding buffer.

### Viral infection and RNA isolation

#### SARS-CoV-2

Viral stocks were propagated as previously reported^87^. Briefly, Vero-TMPRSS2 cells were infected with SARS-CoV-2 Victoria 02/20 strain at a multiplicity of infection (MOI) of 0.003 and incubated for 48–72 h until visible cytopathic effect was observed. Viral titers were then determined by plaque assay from clarified supernatants. For sequencing, Calu-3 cells were infected at an MOI of 1 for 1 h at 37 °C. The inoculum was then removed, cells washed thrice with PBS and incubated in Advanced DMEM with 10% FCS at 37 °C for 24 h. Total RNA was extracted using the RNeasy Mini Kit (Qiagen) and infection confirmed plaque assay and RT-qPCR using primers for the viral N gene: forward 5’-CACATTGGCACCCGCAATC-3’, reverse 5’-GAGGAACGAGAAGAGGCTTG-3’.

#### HCV and ZIKV

As previously reported^88^, ZIKV MP1751 strain was propagated in Vero cells, and concentrated with 8% PEG in NTE buffer. For infection, Huh7.5 were inoculated with either HCV or ZIKV (MOI 1) for 180 min, before extensive washing with PBS. Media was replaced and infected cells were cultured for 72 h before lysing in RLT buffer. RNAs were extracted using RNeasy Mini Kit (Qiagen) and infection confirmed using RT-qPCR quantification of viral RNAs using specific primers pairs. HCV: forward 5’-TCCCGGGAGAGCCATAGTG-3’, reverse 5’-TCCAAGAAAGGACCCAGTC-3’; ZIKV: forward 5’-TCGTTGCCCAACACAAG-3’, reverse 5’-CCACTAATGTTCTTTTGCAGACAT-3’; and RPLP0: forward 5’-GCAATGTTGCCAGTGTCTG-3’, reverse 5’-GCCTTGACCTTTTCAGCAA-3’.

#### HDV

HDV inoculum was prepared by concentrating the culture supernatant of Huh-7 cells transfected with pSVL(D3) and pT7HB2.7 plasmids as previously reported^89^. HepG2-NTCP cells were differentiated with 2.5% DMSO-containing culture medium for 72 h prior to viral infection, before inoculation with HDV (MOI 50) in the presence of 4% PEG8000 and 2.5% DMSO for 24 h. After 24 h, cells were washed with PBS and cultured for an additional 5 days in the presence of 2.5% DMSO. Cells were lysed in RLT buffer, and total cellular RNA was purified using RNeasy Mini Kit (Qiagen) and RNase-free DNase Set (Qiagen). Infection was confirmed by RT-qPCR using specific primers to detect HDV transcripts.

#### SINV

SINV was produced from pT7-SVwt plasmid^90^ that was first linearized with XhoI and purified to use it as a template for in vitro RNA transcription with HiScribe T7 ARCA mRNA kit (NEB). Transcribed viral RNA was transfected into BHK-21 using Lipofectamine 3000 reagent (Invitrogen) according to manufacturer’s instruction. Viruses were collected from the supernatant 24 h later and cleared by centrifugation at 2,000 r.p.m for 5 min followed by filtration with 0.45μm PVDF syringe filter units (Merck) and stored at –80 °C. Cleared supernatants were titrated by plaque assay.

PolyA^+^ RNA purification was performed based on the previously described protocols^91,92^, with the following alterations: A549 cells were seeded in two 10 cm dishes in DMEM 10% FBS 24 h prior to infection to reach 80% confluence. Next, cells were either mock-infected or infected using 0.1 MOI of SINV for 1 h in serum-free DMEM at 37 °C, followed by the replacement of the medium with DMEM supplemented with 2% FBS and incubated for 18 h. Cells were lysed with 1 ml of lysis buffer (20 mM Tris-HCl pH 7.5, 500 mM LiCl, 0.5% (w/v) LiDS, 1 mM EDTA, 0.1% IGEPAL (NP-40) and 5 mM DTT). Lysates were homogenized by passing the lysate at high speed through a 5 ml syringe with a 27G needle, repeating this process until the lysate was fully homogeneous. Then, the whole lysate was incubated with pre-equilibrated oligo(dT)25 magnetic beads (NEB) for 1 h at 4 °C with gentle rotation. Beads were collected in the magnet and washed twice with 2 ml of buffer 1 (20 mM Tris-HCl pH 7.5, 500 mM LiCl, 0.1% (w/v) LiDS, 1 mM EDTA, 0.1% IGEPAL and 5 mM DTT) for 5 min at 4 °C with gentle rotation, followed by two washes with buffer 2 (20 mM Tris-HCl pH 7.5, 500 mM LiCl, 1 mM EDTA, 0.01% IGEPAL and 5 mM DTT). Beads were then washed twice with 2 ml of buffer 3 (20 mM Tris-HCl pH 7.5, 200 mM LiCl, 1 mM EDTA and 5 mM DTT) at room temperature. Finally, beads were resuspended in 50 µl of elution buffer and incubated for 7 min at 55 °C with agitation. Eluates were stored at –80 °C. Approximately 5 µg of polyA^+^ RNAs were used for further rRNA removal using the Ribo-Zero kit from the TruSeq Stranded Total RNA LT Kit (Illumina). Subsequently, RNAclean XP (Beckman Coulter) purification was conducted, and RNA were finally eluted into 10–15 µl of nuclease-free water. The concentration and quality of the RNA were assessed using Qubit and RNA Bioanalyzer.

#### Preparation of IVT RNA control

100 ng polyA^+^ RNA was annealed with 2 μl of 10 μM Oligo(dT)_30_VN primer (5’-TTTTTTTTTTTTTTTTTTTTTTTTTTTTTTVN-3’) and 2 μl of 10 mM dNTP mix (NEB) in 12 μl solution, incubated at 70 °C for 5 min and held at 4 °C. Next, 5 μl 4× Template Switching RT buffer (NEB), 1 µl of 75 μM T7-TSO (5’-/5Biosg/ACTCTAATACGACTCACTATAGGGAGAGGGCrGrGrG-3’), and 2 μl 10× Template Switching RT Enzyme Mix (NEB) were added to the mixture and the reaction was incubated at 42 °C for 90 min followed by 85 °C for 5 min. The second strand synthesis was then performed by adding 100 μl Q5 Hot Start High Fidelity Master Mix (NEB), 10 μl RNase H (NEB), and 70 μl nuclease-free H_2_O to the cDNA mixture and incubation at 37 °C for 15 min, 95 °C for 1 min, and 65 °C for 10 min. The double-stranded cDNA product was purified with 0.8× AMPure XP beads (Beckman Coulter). The IVT RNA control was prepared by T7 *in vitro* transcription using the purified cDNA product and HiScribe T7 High Yield RNA Synthesis Kit according to the manufacturer’s protocol. To remove cDNA template, IVT RNA was then treated with Turbo DNase and purified by Zymo-IC Column with RNA binding buffer.

### BACS for Ψ detection

RNA was fragmented by NEBNext Magnesium RNA Fragmentation Module according to the manufacturer’s protocol. The fragmented RNA was 3’-end repaired using T4 PNK (NEB) and ligated to RNA adapter (5’-/5rApp/AGATCGGAAGAGCGTCGTG/3SpC3/-3’) using T4 RNA Ligase 2, truncated KQ (NEB). Excess adapters were digested using 5’-Deadenylase (NEB) and RecJ_f_ (NEB). For BACS, RNA was treated with 250 mM 2-bromoacrylamide in 625 mM phosphate buffer (pH 8.5) and incubated at 85 °C for 30 min. RNA was reverse transcribed using RT primer (5’-ACACGACGCTCTTCCGATCT-3’) and Maxima H^−^ Reverse Transcriptase (Thermo). Excess RT primers were digested using Exo I (NEB). RNA was hydrolyzed by NaOH (Sigma) and then neutralized by HCl (Sigma). The cDNA was ligated to cDNA adapter (5’-/5Phos/NNNNNNAGATCGGAAGAGCACACGTCTG/3SpC3/-3’) using T4 RNA Ligase 1, high concentration (NEB). The ligated cDNA was amplified with NEBNext Multiplex Oligos for Illumina (96 Unique Dual Index Primer Pairs) and NEBNext Ultra II Q5 Master Mix according to the manufacturer’s protocol. The PCR products were purified with 0.8× AMPure XP beads.

#### Data pre-processing

Raw sequencing reads were processed by Cutadapt (v.4.2)^93^ to remove low-quality bases (-q 20) and short reads (-m 18), as well as to trim adaptors. 6mer UMI were extracted by UMI-tools extract (v.1.0.1)^94^ and used for deduplication. Paired reads were then merged into single reads using fastp (v.1.0.1)^95^.

#### Read alignment

Cleaned reads were first mapped to synthetic spike-ins and rRNA references using bowtie2 (v.2.4.4)^96^. The unaligned reads were subsequently mapped to snoRNA references and then to tRNA references. Human snoRNA sequences that belong to HGNC “Small nucleolar RNAs” gene group were downloaded from RefSeq. Duplicate snoRNA sequences were removed. High-confidence human tRNA sequences (hg38) were downloaded from GtRNAdb^97^. Only non-redundant tRNA sequences were kept and appended with a “3’-CCA” end. Finally, unmapped reads were aligned to human genome (hg38) with GENCODE v.43 annotation by STAR (v.2.7.9a)^98^.

For RNA viruses, the following reference genomes were used: Severe acute respiratory syndrome coronavirus 2 isolate Wuhan-Hu-1 (NC_045512.2), Recombinant Hepatitis C virus J6(5’UTR-NS2)/JFH1 (JF343782.1), Zika virus isolate ZIKV/*H. sapiens*/Brazil/Natal/2015 (NC_035889.1), Hepatitis Delta Virus sequence from the pSVL(D3) plasmid^99^ (Addgene plasmid #29335) (https://www.addgene.org/29335/), and Sindbis virus (NC_001547.1). For EBV samples, reads were aligned to Epstein-Barr virus (EBV) genome, strain B95-8 (V01555.2).

The aligned reads were then filtered and sorted using samtools (v.1.16.1)^100^. Deduplication was performed using UMI-tools dedup (v.1.0.1)^94^. Finally, mutations are counted by samtools mpileup (v.1.16.1)^100^ and cpup (v.0.1.0) (https://github.com/y9c/cpup).

#### Calling Ψ sites

BACS raw conversion rates were calculated as C/(T+C). The Ψ modification levels were calculated using the linear equation: Ψ modification level = (R–F)/(C–F), where R, F, and C indicated raw conversion rates, motif-specific false-positive rates (from NNUNN spike-in for ncRNAs or IVT-BACS library for polyA-tailed RNA), and motif-specific conversion rates (from NNΨNN spike-in), respectively. The binomial test was applied for Ψ calling in ncRNAs using the motif-specific false-positive rate; the contingency table test was applied for Ψ calling in polyA^+^ RNA using the IVT control.

#### Calling A-to-I editing sites

A-to-I editing sites were called based on ICE-seq protocol^101^, with minor modifications.

#### RNA structure visualization

The RNA-RNA interactions were visualized using r2r (v.1.0.6)^102^. The snoRNA-rRNA interactions were adapted from snoRNA Atlas^46^.

#### Downstream analysis

The snoRNA box and guide sequences were downloaded from snoDB 2.0 (ref.^103^). In the metagene analysis, snoRNA sequences that displayed considerable similarity were streamlined, retaining only one representative snoRNA. The annotation of Ψ sites identified in polyA-tailed RNA was performed using bedtools intersect (v.2.30.0)^104^ with GENCODE v.43 annotation. Read counts obtained from featureCounts (v.1.6.4)^105^ were normalized based on sequencing depth and gene length using the transcripts per million (TPM) method. GO analysis was performed with mRNA Ψ sites using enrichR^106^. Sequence logos were generated using ggseqlogo^107^.

#### Published data

Related published data were downloaded from the Gene Expression Omnibus (GEO) database: BID-seq for HeLa cells (GSE179798)^29^. BID-seq data were processed using the original pipeline (BID-seq^29^) and updated pipeline (BID-pipe^31^), respectively.

## Acknowledgements

We would like to acknowledge K. Dunning for editing the manuscript and T. McMahon for helping with UHPLC-MS/MS experiments. We would like to thank C. Rice (Rockefeller University), S. Urban (University of Heidelberg), and N. Zitzmann (University of Oxford), for the generous provision of Huh7.5, HepG2-NTCP, and Calu-3 cell lines, respectively. Furthermore, we would like to acknowledge C. Rice (Rockefeller University), A. Kohl (CVR, University of Glasgow), and W. James (University of Oxford), for the kind provision of HCV J6/JFH1, ZIKV, and SARS-CoV-2 stocks, as well as S. Camille (Université de Tours) for the pT7HB2.7 plasmid. This work was funded by the Ludwig Institute for Cancer Research (to C.-X. S., X. L., and S.K.). C.-X. S. lab is also supported by National Institute for Health Research (NIHR) Oxford Biomedical Research Centre (BRC). J.A.M. lab is funded by Wellcome Investigator Award (200838/Z/16/Z), Wellcome Discovery Award (225198/Z/22/Z), and Chinese Academy of Medical Sciences Innovation Fund for Medical Science, China (2018-I2M-2-002). A.C. is funded by the European Research Council (ERC) Consolidator Grant ‘vRNP-capture’ 101001634 and the MRC grants MR/R021562/1 and MC_UU_00034/2. H.X. and L.K. are supported by China Scholarship Council (CSC). A.E.-B. is funded by Fundación Ramón Areces post-doctoral fellowship program. The views expressed are those of the author(s) and not necessarily those of the NHS, the NIHR, or the Department of Health.

## Author contributions

H.X. and C.-X.S. conceived and designed the study. H.X. performed the experiments with the help from L.K., K.A.M., X.C., A.I., S.K. and X.L. P.A.C.W., J.M.H., S.T. and J.A.M. performed isolation of SARS-CoV-2, HCV, ZIKV and HDV RNA. A.E.-B., G.W. and A.C. performed isolation of SINV RNA. L.K. performed the computational analysis with the help from J.C. H.X., L.K. and C.-X.S. wrote the manuscript.

