## Supplementary Figure 1-15 and Supplementary Table 1-2 for "Absolute quantitative and base-resolution sequencing reveals comprehensive landscape of pseudouridine across the human transcriptome"

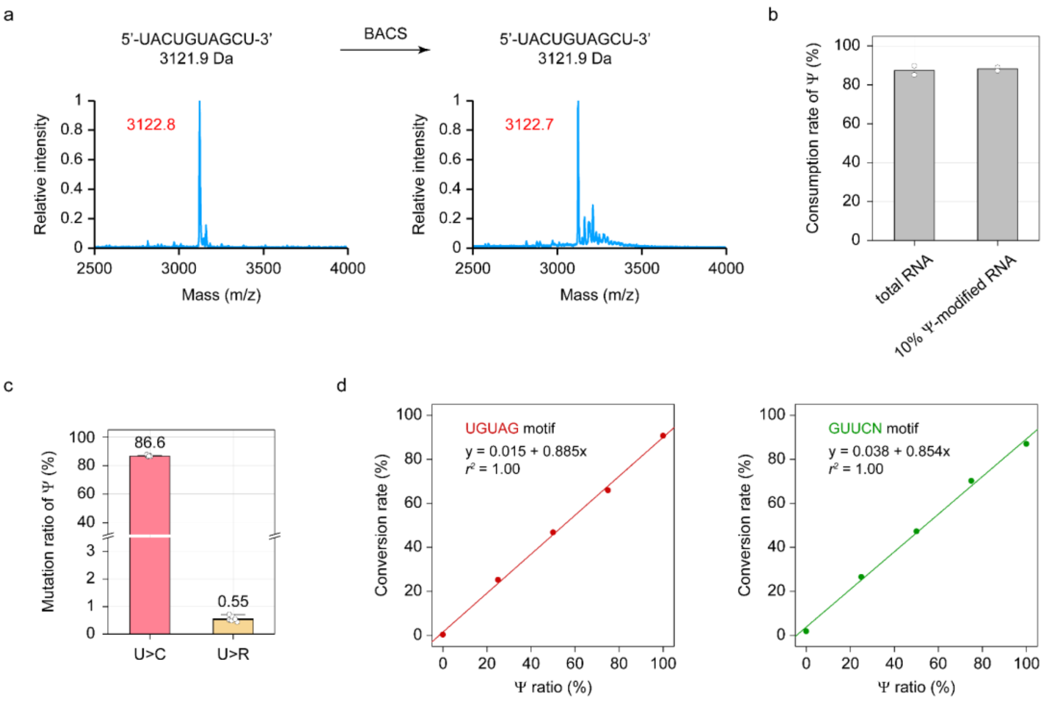


**Supplementary Fig. 1 | Performance of BACS on model RNA. (a)** MALDI characterization of BACS labeling of a 10mer unmodified RNA oligonucleotide. Calculated mass is shown in black. Observed mass is shown in red. Experiment was performed once. **(b)** Consumption rates of Ψ in HeLa total RNA and 1.8-kb 10% Ψ-modified RNA upon BACS treatment, quantified by UHPLC-MS/MS. Data are presented as means of two independent experiments. **(c)** Mutation ratios of Ψ sites in 72mer model RNA after BACS treatment. Data are shown as means ± s.d. of six independent experiments ($n$ = 6). **(d)** BACS calibration curve for quantification of Ψ stoichiometry in UGUAG (red) and GUUCN (green) motif. Experiment was performed once.

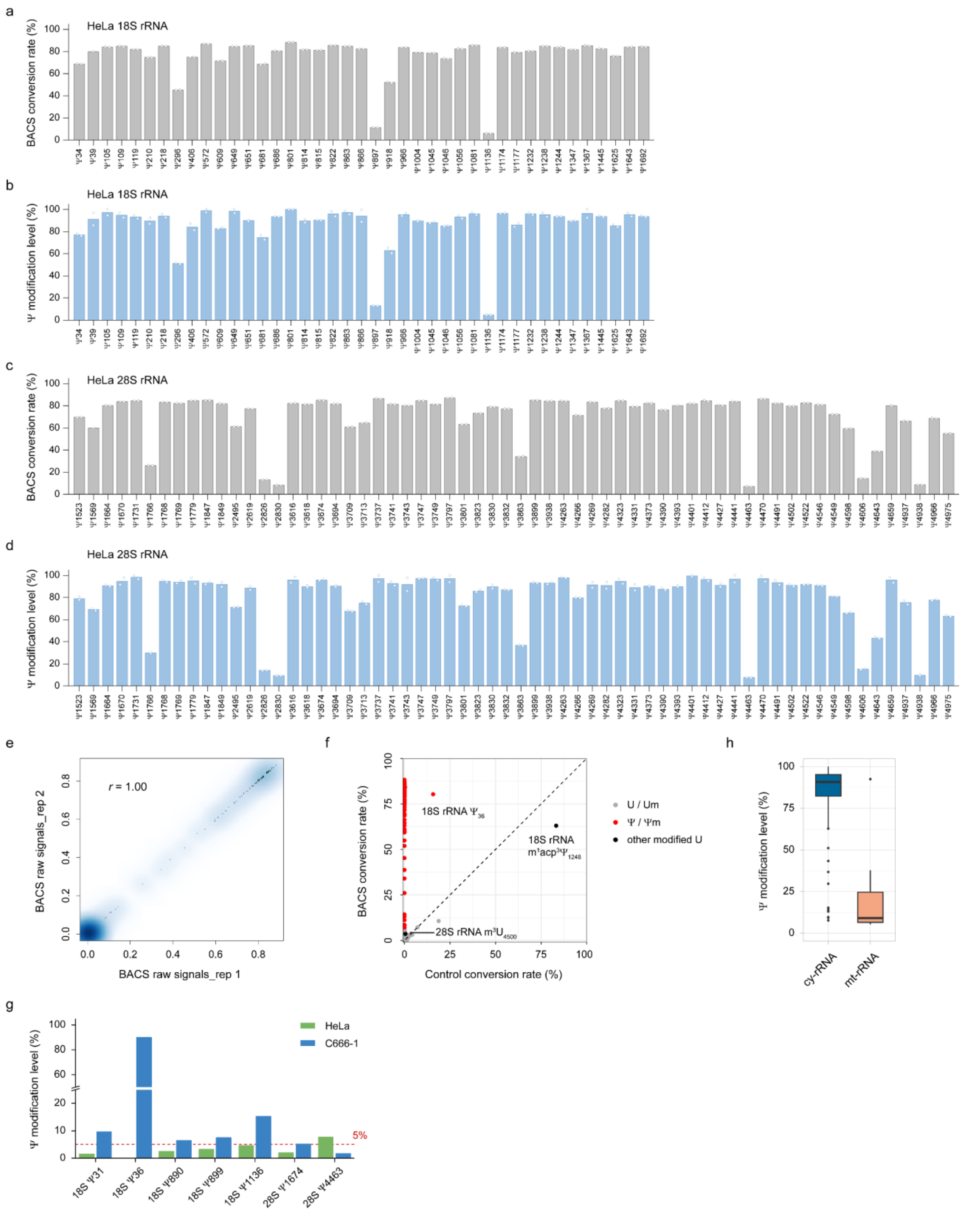


**Supplementary Fig. 2 | BACS validated known Ψ sites in human rRNA. (a)** BACS conversion rates of Ψ sites identified in HeLa 18S rRNA. Data are presented as means of two independent experiments. **(b)** Modification levels of Ψ sites detected in HeLa 18S rRNA. Data are presented as means of two independent experiments. **(c)** BACS conversion rates of Ψ sites identified in HeLa 28S rRNA. Data are presented as means of two independent experiments. **(d)** Modification levels of Ψ sites detected in HeLa 28S rRNA. Data are presented as means of two independent experiments. **(e)** Correlation density plot between two biological replicates of BACS. The color scale represents density. **(f)** Comparison of the conversion rates in HeLa cy-rRNAs between BACS and control samples. **(g)** Comparison of the modification levels of selected Ψ sites in cy-rRNAs between HeLa (green) and C666-1 (blue) cell lines. **(h)** Comparison of the modification levels of Ψ sites in HeLa cy-rRNAs and mt-rRNAs. Boxplots indicate medians, quantiles, extreme values, and outliers (cy-rRNA, $n$ = 104; mt-rRNA, $n$ = 9).

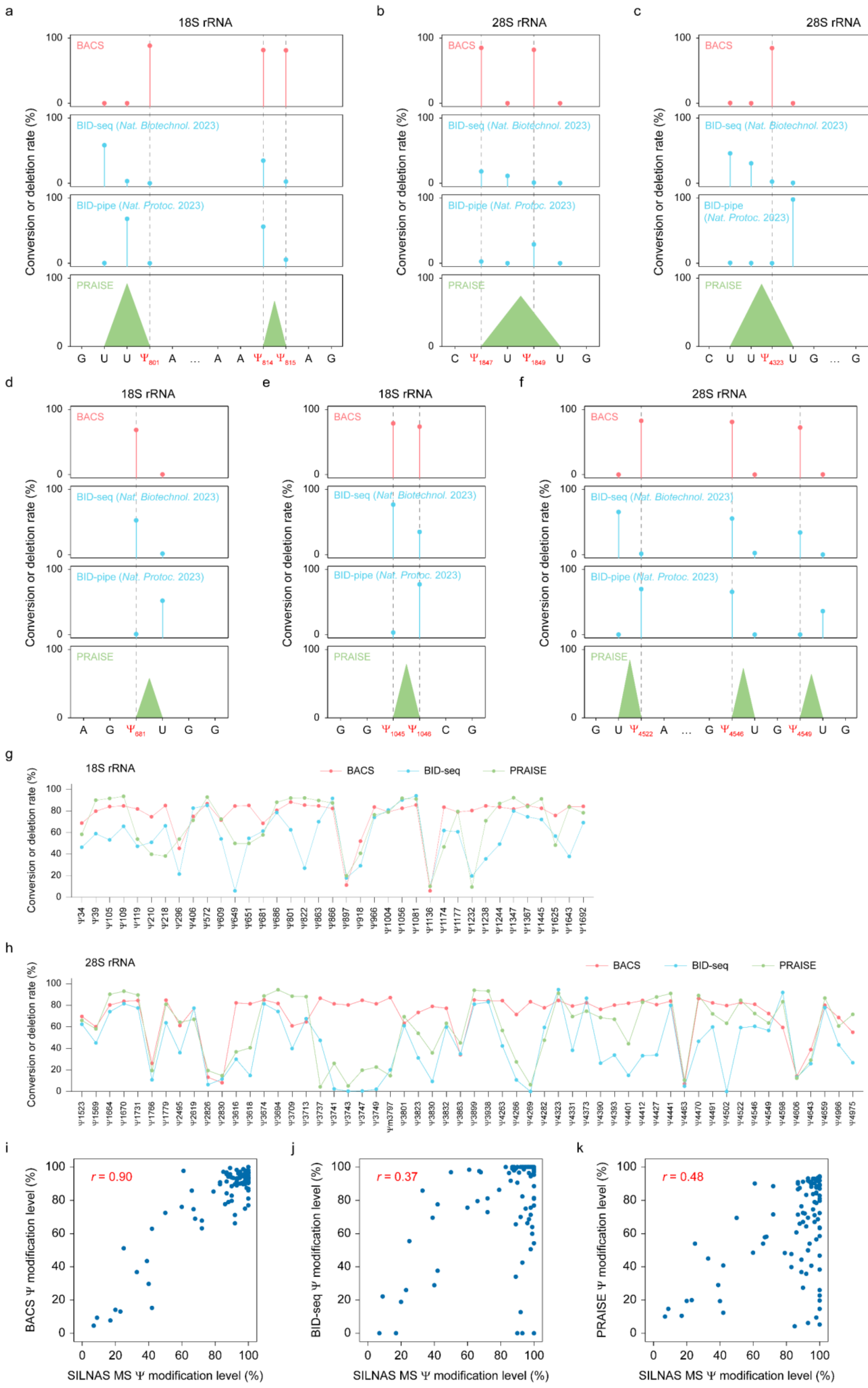

a
18S rRNA
b
28S rRNA
c
28S rRNA
BACS
BID-seq (Nat. Biotechnol. 2023)
BID-pipe (Nat. Protoc. 2023)
PRAISE
Conversion or deletion rate (%)
d
18S rRNA
e
18S rRNA
f
28S rRNA
g
18S rRNA
BACS
BID-seq
PRAISE
h
28S rRNA
i
r = 0.90
BACS Ψ modification level (%)
SILNAS MS Ψ modification level (%)
j
r = 0.37
BID-seq Ψ modification level (%)
SILNAS MS Ψ modification level (%)
k
r = 0.48
PRAISE Ψ modification level (%)
SILNAS MS Ψ modification level (%)

**Supplementary Fig. 3 | Comparison of BACS and BS-based methods for Ψ detection in human cy-rRNAs. (a–f)** Examples of BACS, BID-seq, BID-pipe, and PRAISE results in selected consecutive uridine regions of 18S and 28S rRNA. Due to their deletion signals, BS-based methods cannot determine the exact position or identify the number of Ψ within consecutive uridine contexts. By default, the aligner will position the deletion at the 5'-most uridine, as shown in the BID-seq panel. After realignment, this issue cannot be fully resolved, as shown in the BID-pipe panel. PRAISE considers consecutive uridines as a whole for Ψ calling, resulting in broad peak signals. **(g,h)** Comparison of the conversion rates of BACS (pink) with the deletion rates of BID-seq (blue) and PRAISE (green) for selected Ψ sites in 18S rRNA **(g)** and 28S rRNA **(h)**. Because BID-seq and PRAISE cannot quantify multiple Ψ sites (≥ 2) located in the same consecutive uridine context, these sites were excluded. **(i–k)** Correlation of cy-rRNA Ψ modification levels reported by SILNAS MS with those reported by BACS **(i)**, BID-seq **(j)**, and PRAISE **(k)**.

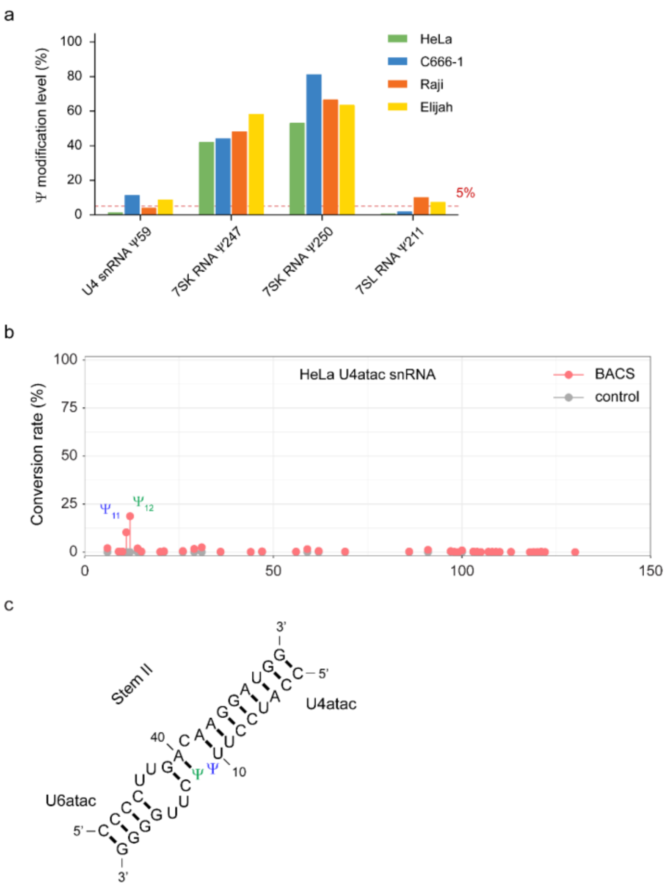


**Supplementary Fig. 4 | BACS identified conserved and novel Ψ sites in human spliceosomal snRNA. (a)** Comparison of the modification levels of selected Ψ sites in U4 snRNA, 7SK RNA, and 7SL RNA across HeLa (green), C666-1 (blue), Raji (orange), and Elijah (yellow) cell lines. **(b)** Conversion rates of BACS (pink) and control (grey) samples in U4atac snRNA, showing the novel (blue) and known (green) Ψ site. Data are presented as means of two independent experiments. **(c)** Base pairing interactions between U4atac and U6atac snRNAs in stem II region. Blue and green color denote the novel $\Psi_{11}$ and known $\Psi_{12}$ site, respectively.

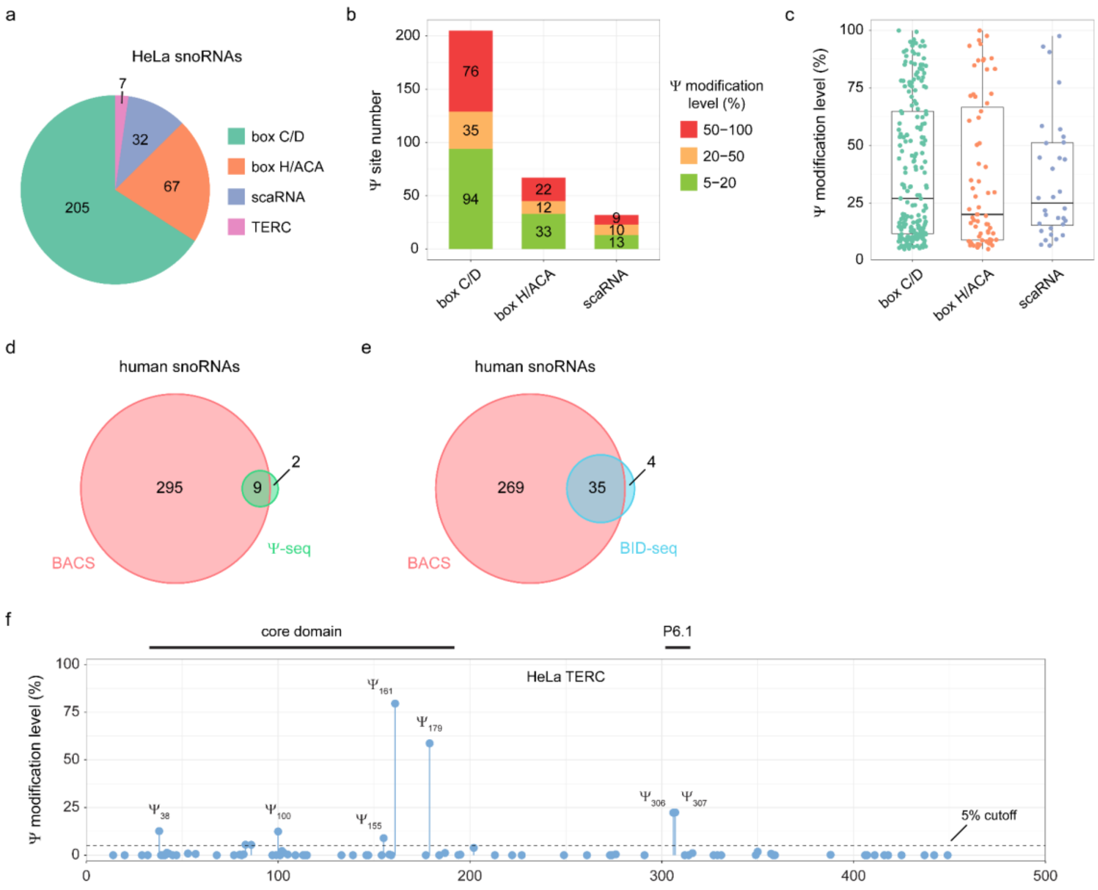


**Supplementary Fig. 5 | BACS detected abundant Ψ sites in human snoRNAs. (a)** Numbers of Ψ sites identified in HeLa snoRNAs and TERC. **(b)** Numbers of Ψ sites with high (50–100%, red), medium (20–50%, yellow), and low (5–20%, green) modification levels identified in HeLa box C/D snoRNAs, box H/ACA snoRNAs, and scaRNAs. **(c)** Modification level distributions of Ψ sites in HeLa box C/D snoRNAs, box H/ACA snoRNAs, and scaRNAs. Boxplots visualize all Ψ sites in each class of snoRNAs, indicating medians, quantiles, and extreme values (box C/D, $n$ = 205; box H/ACA, $n$ = 67; scaRNA, $n$ = 32). **(d)** Venn diagram illustrating the overlap of Ψ sites detected in human snoRNAs between BACS and Ψ-seq. **(e)** Venn diagram illustrating the overlap of Ψ sites detected in human snoRNAs between BACS and BID-seq. **(f)** Ψ modification levels in HeLa TERC, with each identified Ψ site labeled accordingly. Data are presented as means of two independent experiments.

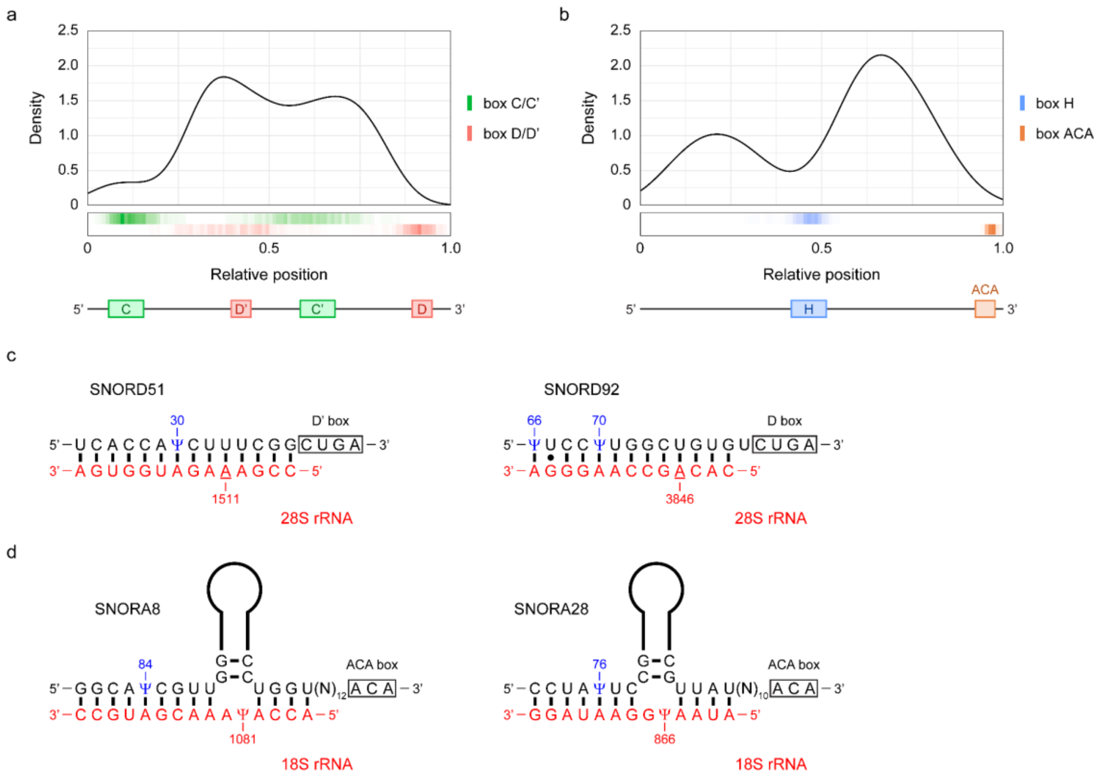


**Supplementary Fig. 6 | Potential involvement of Ψ in regulating the guiding activity of human snoRNAs. (a)** Metagene profile of Ψ sites in box C/D snoRNAs. Red and green color denote the box C/C' and D/D', respectively. **(b)** Metagene profile of Ψ sites in box H/ACA snoRNAs. Orange and blue color denote the box H and ACA, respectively. **(c,d)** Potential base pairing interactions between snoRNAs (black) and their targets in rRNA (red): **c.** box C/D snoRNAs and **d.** box H/ACA snoRNAs. Identified snoRNA Ψ sites are highlighted in blue. 2'-*O*-methylation targets in rRNA are underlined. Structures are adapted from snoRNA Atlas[1].

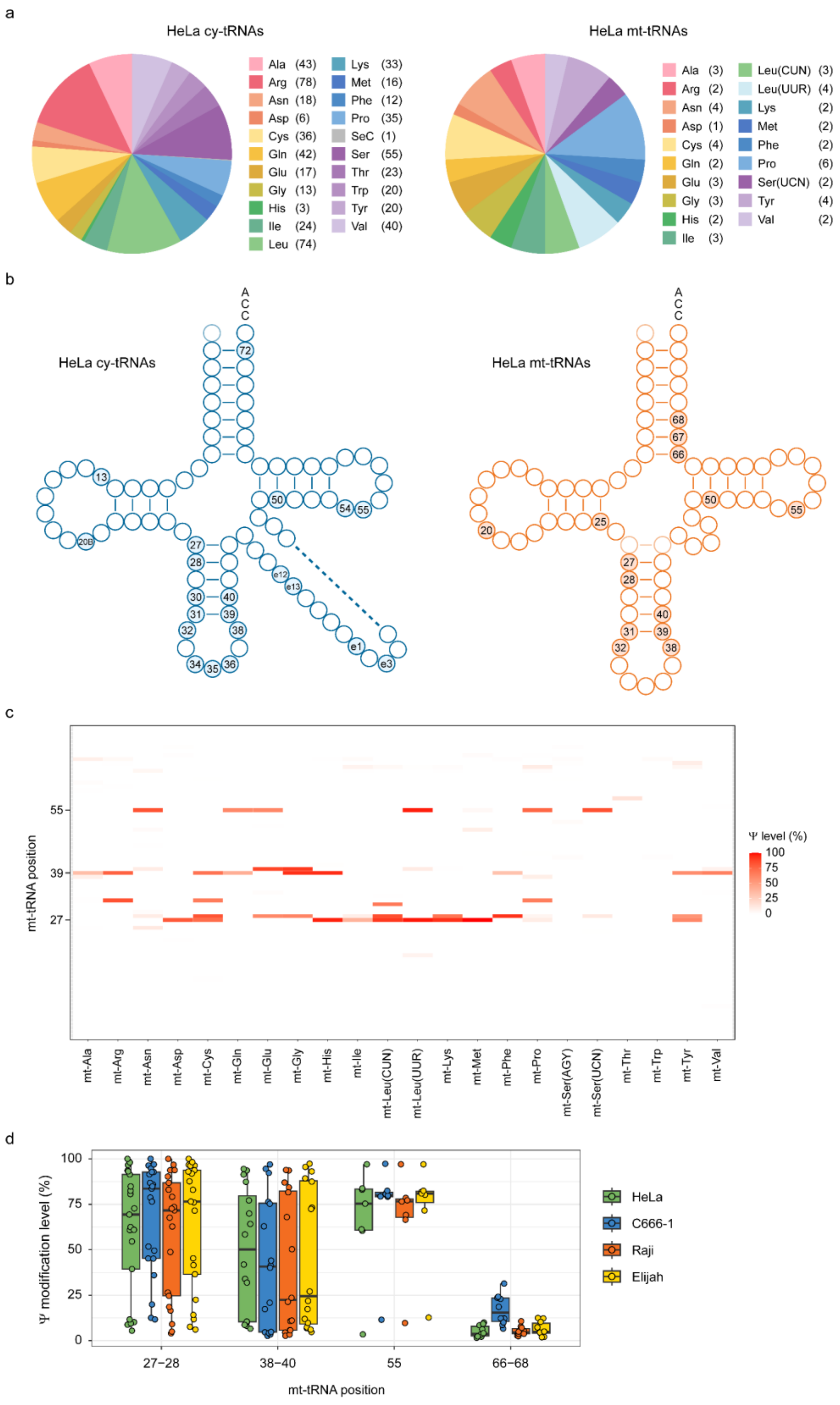


**Supplementary Fig. 7 | BACS revealed a comprehensive map of Ψ in human tRNA. (a)** Distributions of Ψ sites identified in each cy-tRNA (left) and mt-tRNA (right) isotype from HeLa cells. **(b)** Integrated view of the Ψ profiles of HeLa cy-tRNAs (left) and mt-tRNAs (right). **(c)** Heatmap of the Ψ modification levels in HeLa mt-tRNAs. **(d)** Comparison of the modification levels of Ψ sites at selected positions of mt-tRNAs across HeLa (green), C666-1 (blue), Raji (orange), and Elijah (yellow) cell lines. Boxplots visualize all Ψ sites at each position, indicating medians, quantiles, and extreme values (tRNA position: 27–28, *n* = 21; 39–40, *n* = 16; 55, *n* = 7; 66–68, *n* = 10).

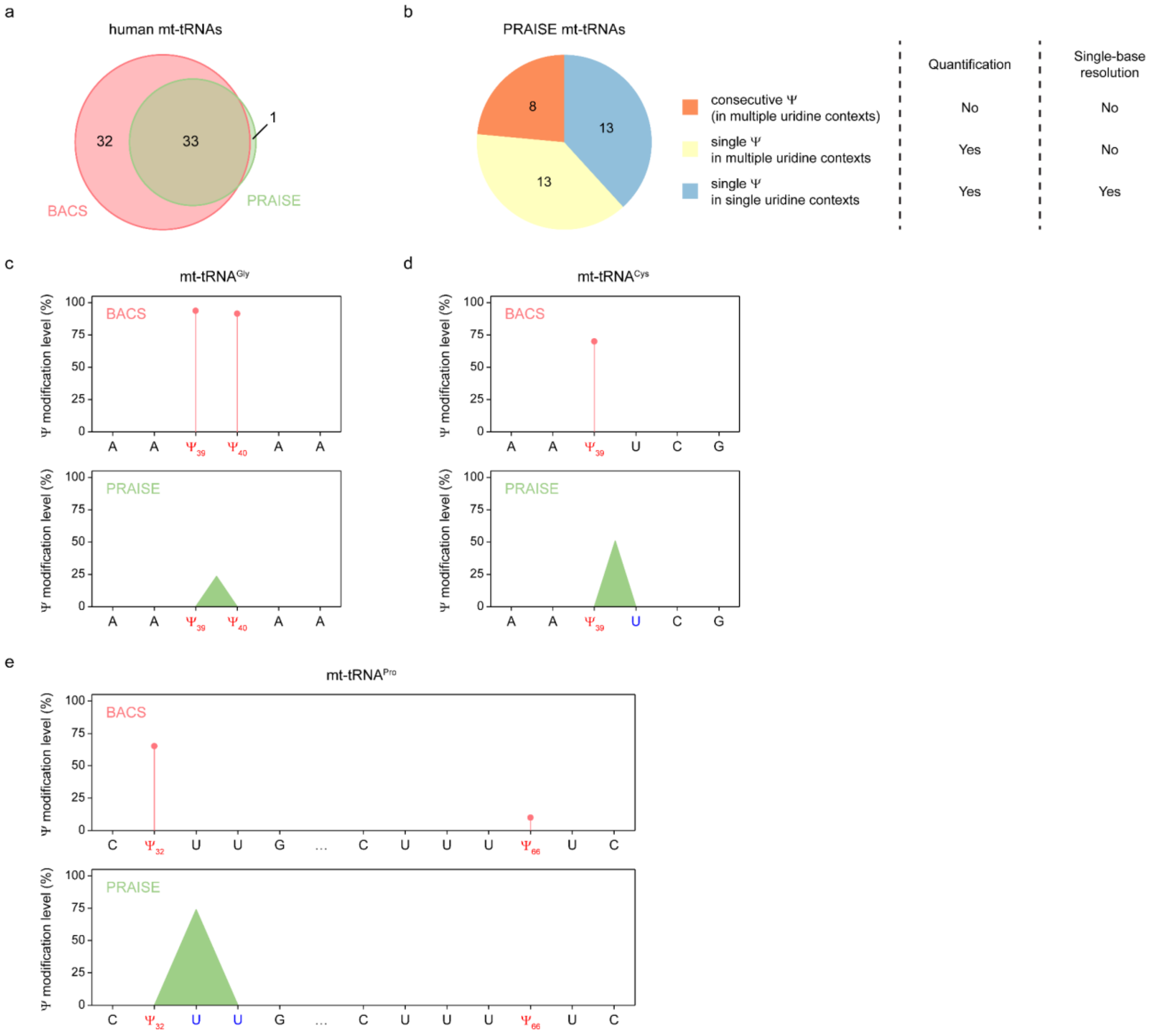


**Supplementary Fig. 8 | Comparison of BACS and PRAISE for Ψ detection in human mt-tRNAs. (a)** Venn diagram illustrating the overlap of Ψ sites detected in human mt-tRNAs between BACS and PRAISE. **(b)** Distribution of mt-tRNA Ψ sites identified by PRAISE. Only single Ψ site in single uridine contexts can be quantitatively identified by PRAISE at single-base resolution. **(c–e)** Examples of BACS and PRAISE results in selected consecutive uridine regions of human mt-tRNAs. Due to their deletion signals, BS-based methods cannot determine the exact position or identify the number of Ψ within consecutive uridine contexts. PRAISE considers consecutive uridines as a whole for Ψ calling, resulting in broad peak signals.

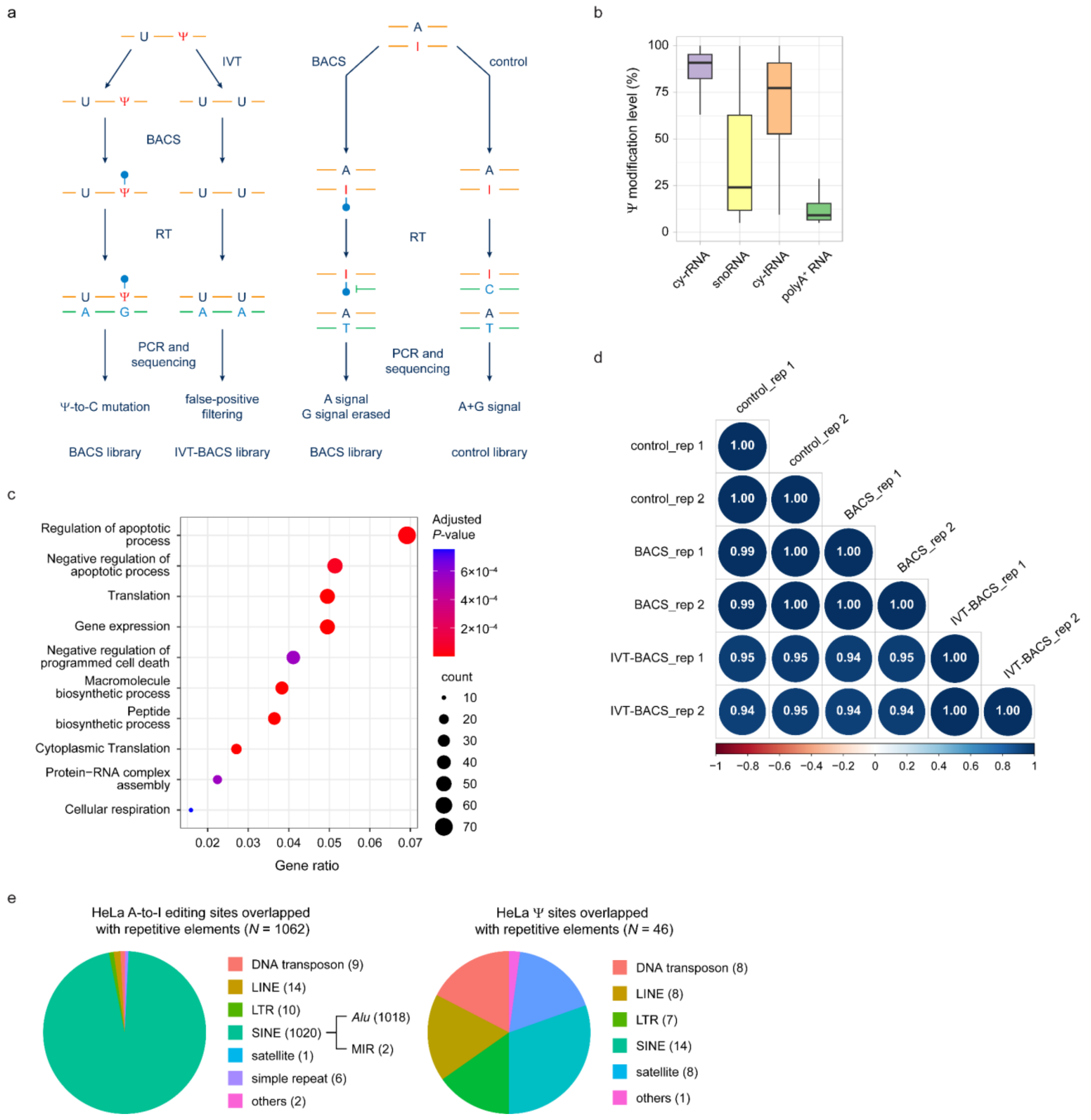


**Supplementary Fig. 9 | Analysis of BACS libraries for polyA-tailed RNA. (a)** Schematic overview of Ψ and A-to-I editing site identification in polyA-tailed RNA. Ψ site calling is based on BACS and IVT-BACS libraries. A-to-I editing site calling is based on BACS and control libraries. **(b)** Comparison of the Ψ modification levels in different RNA species. Boxplots indicate medians, quantiles, and extreme values (cy-rRNA, $n$ = 104; snoRNA, $n$ = 304; cy-tRNA, $n$ = 609; polyA-tailed RNA, $n$ = 1335). **(c)** Gene ontology enrichment analysis (biological process) for HeLa mRNA Ψ sites. **(d)** Correlation of HeLa RNA expression levels between BACS, control, and IVT-BACS libraries. Pearson's $r$ values are shown. **(e)** Distribution of HeLa A-to-I editing and Ψ sites overlapped with repetitive elements.

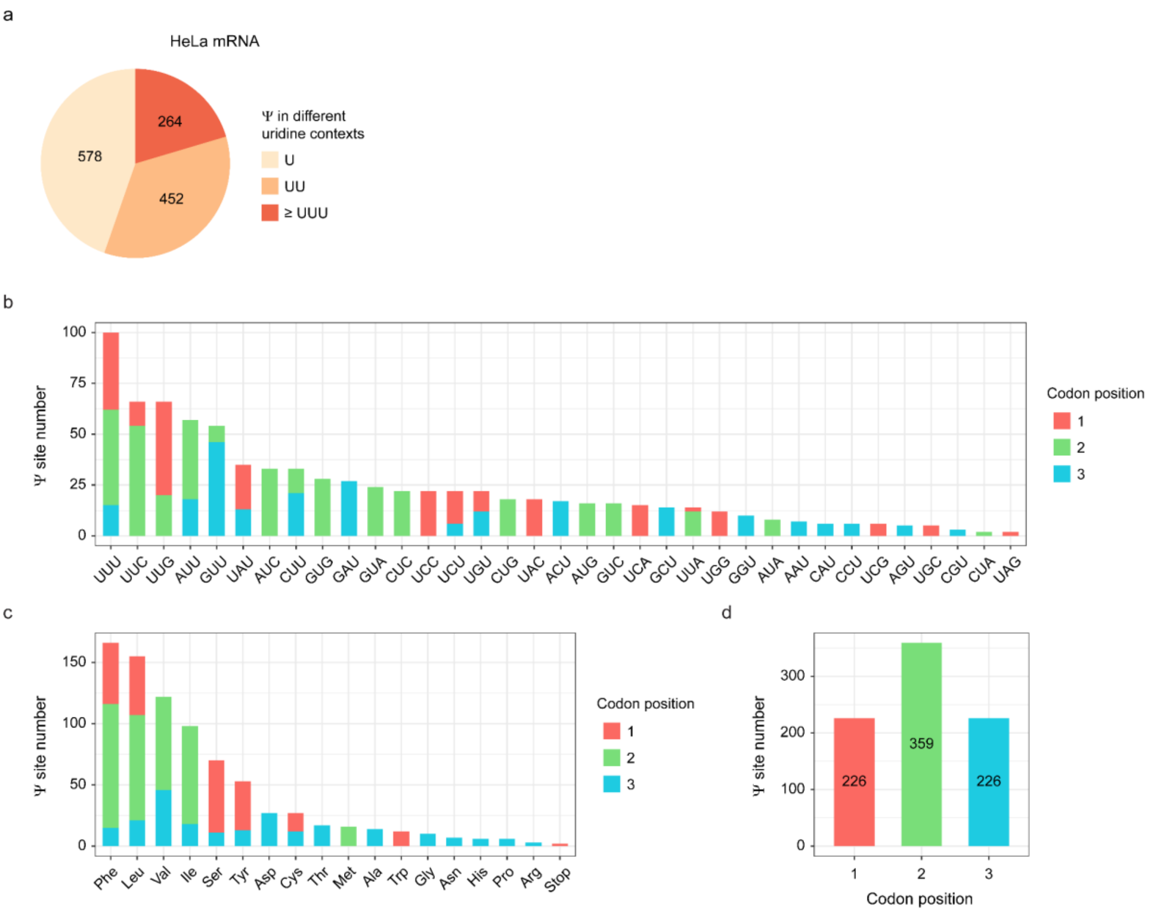


**Supplementary Fig. 10 | Sequence context and codon preference of Ψ in HeLa mRNA. (a)** Distribution of mRNA Ψ sites within single and consecutive uridine contexts. **(b,c)** Numbers of mRNA Ψ sites located in different codons **(b)** and codons encoding different amino acids **(c)**. Red, green, and blue color denote the first, second, and third codon position, respectively. **(d)** Numbers of mRNA Ψ sites located in different codon positions.

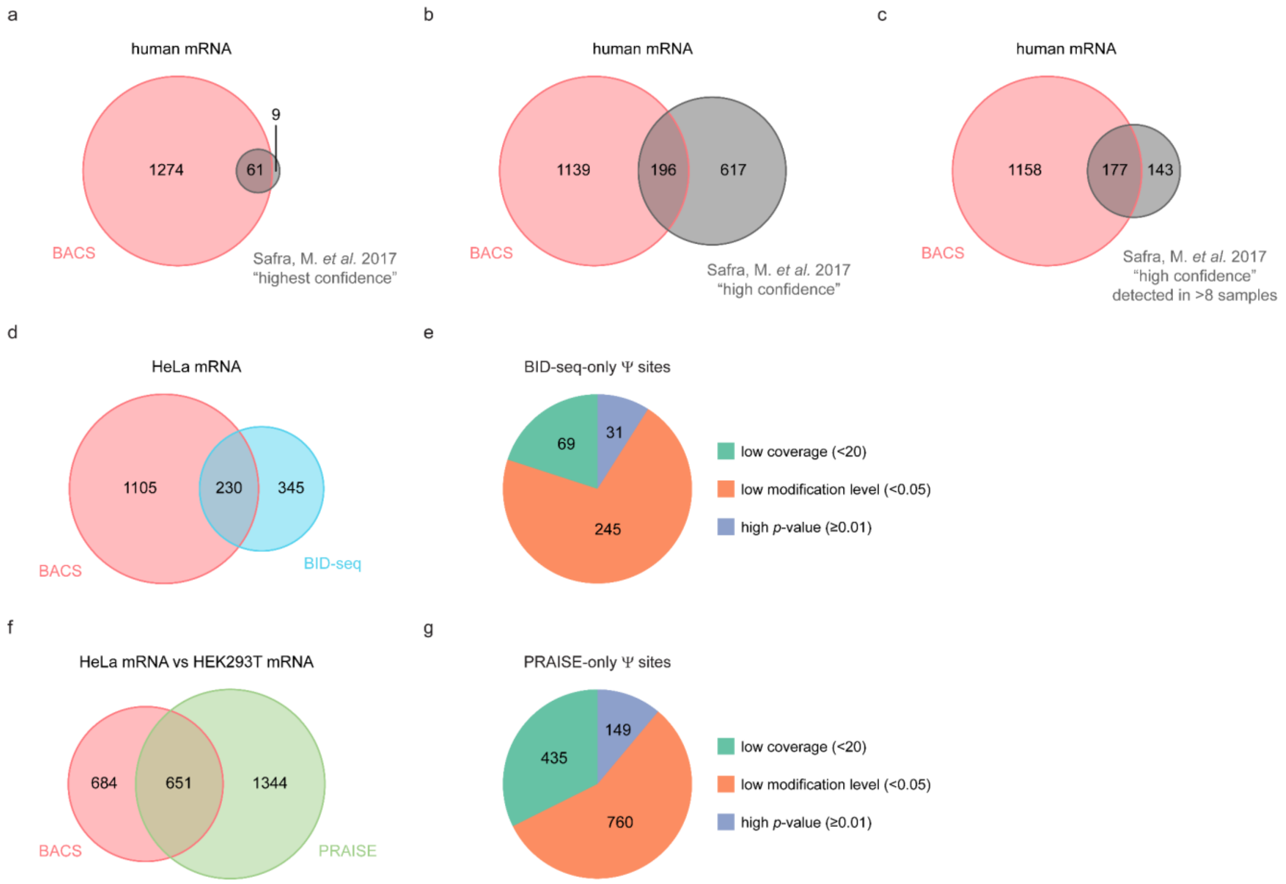


**Supplementary Fig. 11 | Comparison of mRNA Ψ sites identified by BACS with published datasets. (a)** Venn diagram illustrating the overlap of mRNA Ψ sites between BACS and the "highest confidence" list in a consolidated CMC-based dataset[2]. **(b)** Venn diagram illustrating the overlap of mRNA Ψ sites between BACS and the "high confidence" list in a consolidated CMC-based dataset[2]. **(c)** Venn diagram illustrating the overlap of mRNA Ψ sites between BACS and the "high confidence" list in a consolidated CMC-based dataset[2]. Only Ψ sites consistently detected across more than 8 samples in the "high confidence" list were considered. **(d)** Venn diagram illustrating the overlap of mRNA Ψ sites between BACS and BID-seq. **(e)** Distribution of BID-seq-only Ψ sites in BACS dataset. **(f)** Venn diagram illustrating the overlap of mRNA Ψ sites between BACS and PRAISE. **(g)** Distribution of PRAISE-only Ψ sites in BACS dataset.

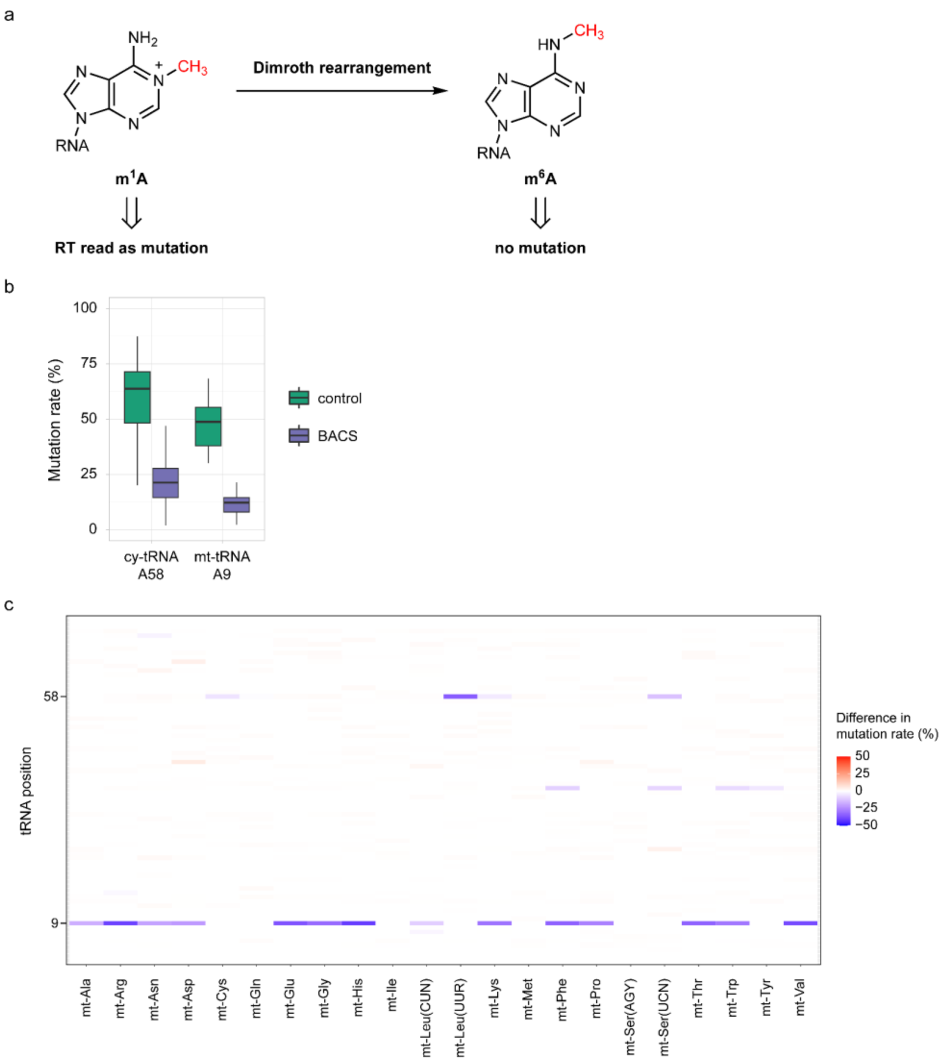


**Supplementary Fig. 12 | BACS enabled simultaneous detection of m$^1$A with Ψ. (a)** Schematic overview of Dimroth rearrangement of m$^1$A to m$^6$A. **(b)** Comparison of the mutation rates of known m$^1$A sites in control (green) and BACS (purple) samples. Boxplots indicate medians, quantiles, and extreme values (cy-tRNA A58, *n* = 170; mt-tRNA A9, *n* = 14). **(c)** Heatmap showing the changes in mutation rates of all adenosine sites in human mt-tRNAs upon BACS treatment. Red and blue color indicate an increase and decrease of mutation rates, respectively.

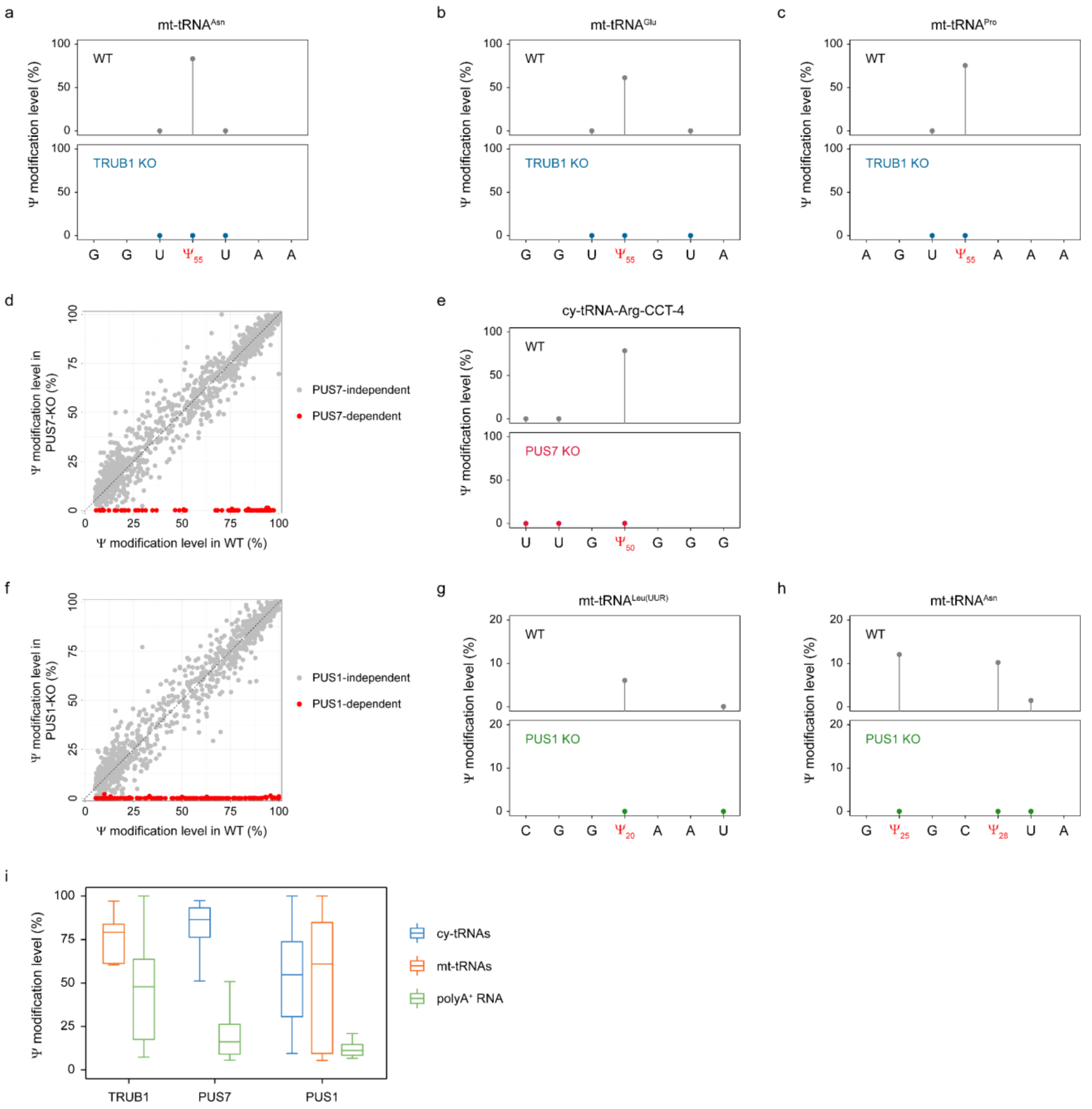


**Supplementary Fig. 13 | BACS assigned responsible PUS enzymes for Ψ sites in the HeLa transcriptome. (a–c)** Comparison of the modification levels of Ψ55 in mt-tRNA$^{Asn}$ **(a)**, mt-tRNA$^{Glu}$ **(b)**, and mt-tRNA$^{Pro}$ **(c)** between WT and TRUB1-KO cell lines. **(d)** Scatter plot illustrating all PUS7-dependant Ψ sites across the HeLa transcriptome. **(e)** Comparison of the modification levels of Ψ50 in cy-tRNA$^{Arg(CCT)}$ between WT and PUS7-KO cell lines. **(f)** Scatter plot illustrating all PUS1-dependant Ψ sites across the HeLa transcriptome. **(g,h)** Comparison of the modification levels of Ψ20 in mt-tRNA$^{Leu(UUR)}$ **(g)** and Ψ25 in mt-tRNA$^{Asn}$ **(h)** between WT and PUS1-KO cell lines. **(i)** Comparison of the modification levels of PUS-dependent Ψ sites across cy-tRNAs (blue), mt-tRNAs (orange), and polyA-tailed RNA (green). Boxplots indicate medians, quantiles, and extreme values (TRUB1: mt-tRNAs, $n$ = 6; polyA-tailed RNA, $n$ = 41; PUS7: cy-tRNAs, $n$ = 69; polyA-tailed RNA, $n$ = 22; PUS1: cy-tRNAs, $n$ = 130; mt-tRNAs, $n$ = 27; polyA-tailed RNA, $n$ = 8).

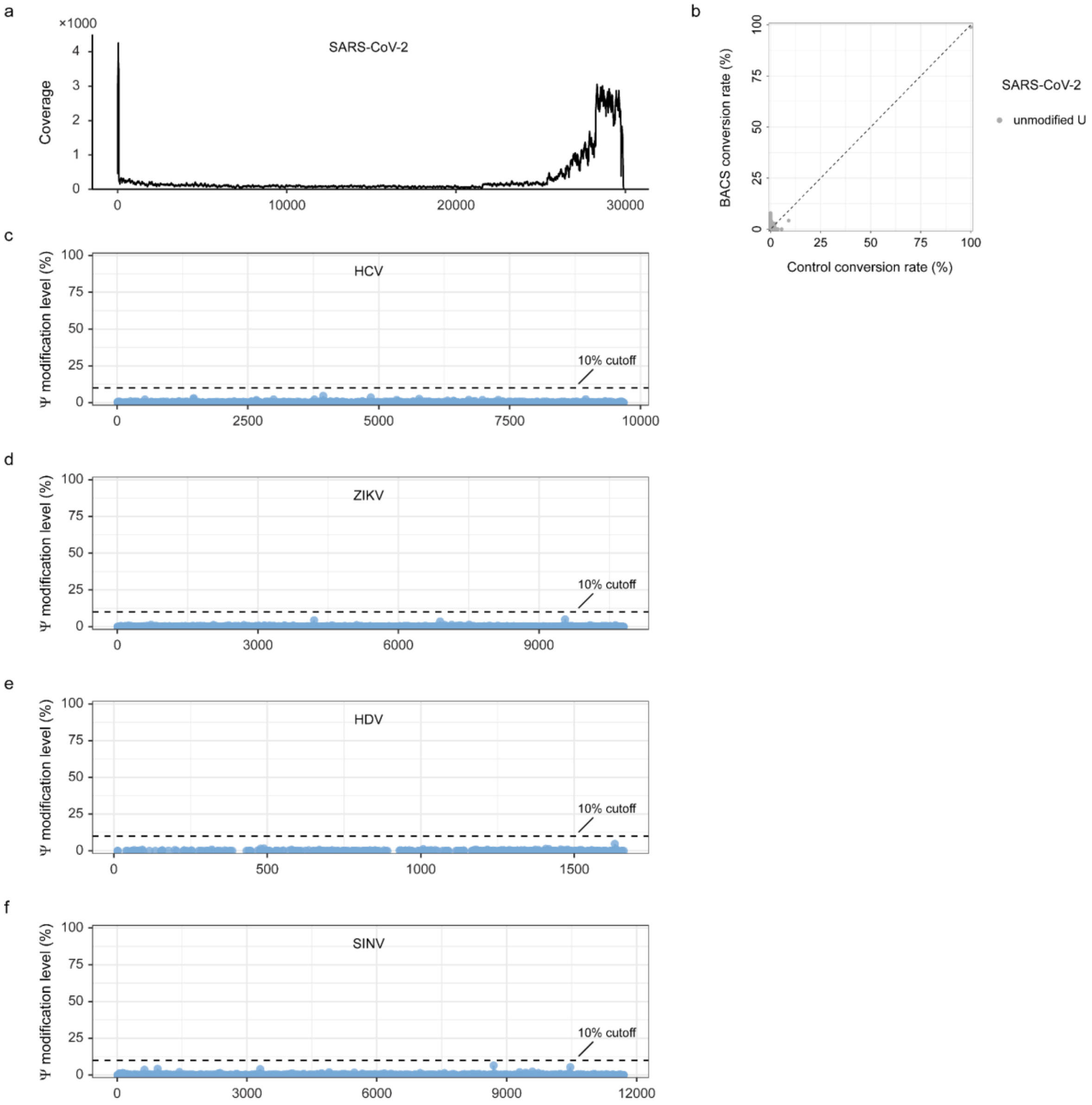


**Supplementary Fig. 14 | Absence of Ψ in transcripts and genomes of RNA viruses. (a)** Coverage of SARS-CoV-2 viral RNA. **(b)** Comparison of the conversion rates in SARS-CoV-2 viral RNA between BACS and control samples. **(c–f)** Ψ modification levels in HCV **(c)**, ZIKV **(d)**, HDV **(e)**, and SINV **(f)** viral RNAs.

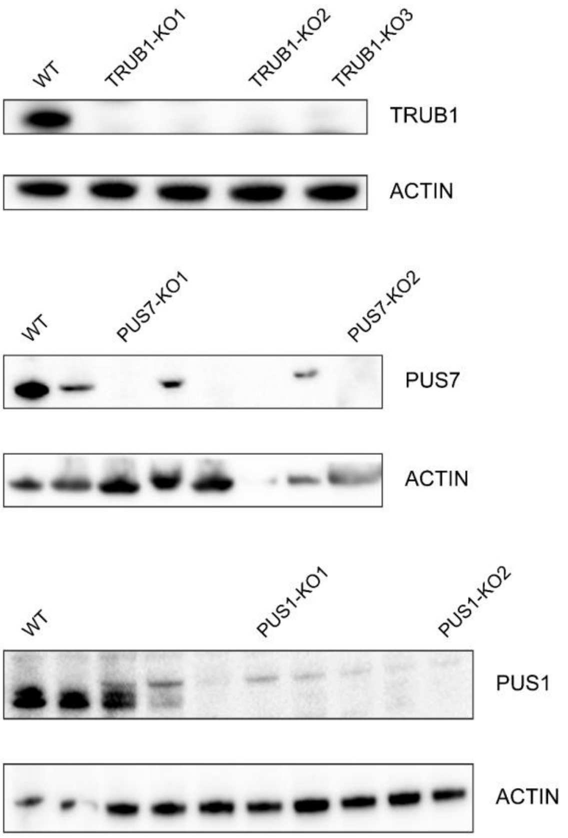


**Supplementary Fig. 15 | Validation of TRUB1-KO, PUS7-KO, and PUS1-KO HeLa cell lines.** β-actin was used as a loading control.

**Supplementary Table 1.** RNA oligonucleotide sequences in this work.

| Name | Sequence (5' to 3') | Source |
|---|---|---|
| for MALDI | | |
| 10mer U-ORN | UACUG**U**AGCU | IDT |
| 10mer Ψ-ORN | UACUG**Ψ**AGCU | IDT |
| for mutation analysis | | |
| 72mer Ψ-ORN | GGGAGAACACACCACAACGAAACCAACGG**Ψ**ACAACAACAGAAA**Ψ**CGAGGACCGAAGCGAAGGCAAAGACAAC | *in vitro* transcription<br>Pseudo-UTP, ATP, CTP, GTP |
| for UHPLC-MS/MS | | |
| 1.8-kb 10% Ψ-modified RNA | T7 *in vitro* transcription using linearized Fluc plasmid (NEB) as template | *in vitro* transcription<br>10% Pseudo-UTP, 90% UTP, ATP, CTP, GTP |
| spike-ins | | |
| 30mer NNUNN | AUGUCUCGACGU**NNUNN**GUUACAGUACCGU | IDT |
| 30mer NNΨNN | GCUUCAAGUUGA**NNΨNN**CAUCGCAAGUGCA | IDT |

**Supplementary Table 2.** Compound-dependent UHPLC-MS/MS parameters used for nucleoside quantification. All the nucleosides were analyzed in the positive mode.

| Compound | Precursor Ion (m/z) | Product Ion (m/z) | RT (min) | Delta RT (min) | CE (V) |
|---|---|---|---|---|---|
| Ψ+H | 245 | 125 | 1.8 | 2.0 | 10.0 |
| rC+H | 244 | 112 | 2.3 | 2.0 | 10.0 |
| rC+Na | 266 | 134 | 2.3 | 2.0 | 10.0 |
| rU+H | 245 | 113 | 3.3 | 2.0 | 10.0 |
| rG+H | 284 | 152 | 8.1 | 2.0 | 10.0 |
| rA+H | 268 | 136 | 12.9 | 2.0 | 8.0 |

RT: retention time; CE: collision energy.
